# Molecular dissection of the ChAHP complex subunits Adnp and Chd4 reveals common and divergent functions in retinal development

**DOI:** 10.64898/2026.10.29.762969

**Authors:** Suma Medisetti, Alena Kalinina, Sujay Shah, Mohammad Amin Fayaz, Samuel Clémot-Dupont, David J Picketts, Pierre Mattar

## Abstract

Epigenetic remodelling is thought to regulate the progression of competence states in neural progenitor cells. Here, we examined the functions of the ChAHP chromatin remodelling complex during retinal development. ChAHP consists of the ATP-dependent nucleosome remodeller CHD4, the zinc finger and homeodomain transcription factor ADNP, and the HP1 heterochromatin proteins (Cbx1/3/5). Notably, both *CHD4* and *ADNP* are genetically associated with neurodevelopmental disorders (NDDs), but ChAHP functions in neural development are not fully understood. In the retina, we found that both ADNP and CHD4 proteins were co-expressed ubiquitously in each cell type throughout development, and the two proteins formed a complex *in vivo*. Next, we generated retina-specific conditional knockouts (cKOs) to both *Adnp* and *Chd4* using the *Chx10-Cre* driver. We observed phenotypic correspondence and divergence. Developmentally, we observed elevated cell death in both *Adnp* and *Chd4* cKOs, but *Adnp* cKOs also exhibited deficits in progenitor numbers and proliferation. As a result, mature *Adnp* cKOs were hypoplastic with reductions in each cell type. We previously described a more complex phenotype in *Chd4* cKOs, where cell fates were imbalanced and progenitor proliferation was prolonged, but these features were absent from the *Adnp* cKO. To understand how ChAHP regulates gene expression, we compared *Adnp* and *Chd4* cKOs via scRNA-seq, which revealed common transcriptional signatures. However, *Chd4* cKOs exhibited more severe effects on gene expression in line with its stronger effects on cell fate. We then performed cut&run-seq, which suggested that Adnp and Chd4 are recruited to the same genomic loci – many of which are also occupied by the Ctcf transcription factor. Taken together, our data suggest that ChAHP regulates retinal growth, but that CHD4 has additional ChAHP-independent functions in regulating progenitor fate decisions. These findings point to the idea that during neurogenesis, some brain regions may have greater dependence on ChAHP, and therefore greater vulnerability to genetic mutation.

**Summary Statement:** To investigate epigenetic processes in retinal development, we conditionally ablated the Adnp and Chd4 subunits of the ChAHP chromatin remodelling complex. Genomic, transcriptomic, and histological analyses reveal that Adnp cooperates with the nucleosome remodeller Chd4 to regulate retinal growth.

## Introduction

During development, progenitor cells in most brain regions produce different types of neurons and glia in chronologically stereotyped sequences. To do this, progenitors undergo transitions in *competence*, meaning that the potential to generate particular neuronal or glial cell fates is sequentially gained and lost over time. Competence transitions are thought to be particularly important in particular brain regions such as the neocortex and retina, where different types of neurons and glia are produced in specific developmental windows. For example, in the retina, 6 types (and as many as ∼120 subtypes) of neurons and glia are generated by a single pool of multipotent retinal progenitor cells (RPCs). In mice, retinal ganglion cells are the first neurons to be produced at the onset of neurogenesis beginning at approximately embryonic day (E) 11.5. Next, RPCs begin to produce other early-born fates, including cone photoreceptors, horizontal cells, and some amacrine subtypes. As the production of these cell types crests, rods begin to be produced. After birth, RPCs lose the competence to produce early-born fates, and become restricted to produce rods, bipolars, certain amacrine subtypes, and Müller glia (Alexiades and Cepko, 1997; Carter-Dawson and LaVail, 1979; Young, 1985). Rod production is particularly dominant postnatally – ultimately accounting for ∼80% of the total cell production (Carter-Dawson and LaVail, 1979; Jeon et al., 1998)..

Cell-intrinsic epigenetic processes have long been thought to be important for regulating competence transitions. Chromatin remodellers such as the polycomb repressive complex have previously been shown to regulate developmental timing (Hirabayashi et al., 2009; Morimoto-Suzki et al., 2014; Tsuboi et al., 2018), including during retinal neurogenesis (Aldiri et al., 2013; Iida et al., 2015; Zhang et al., 2023; Zhang et al., 2015). Moreover, single-cell multi-omic studies confirm that temporal competence transitions correlate with major changes in genome accessibility (Lyu et al., 2021), suggesting that nucleosome remodelling might play a central role. Accordingly, in our prior work, we identified the ATP-dependent nucleosome remodelling enzyme Chd4 (Mi-2β) as a crucial regulator of retinal neurogenesis (Shah et al., 2025). Chd4 was required to schedule the termination of progenitor proliferation at the end of neurogenesis, and also played an important role in balancing cell type production and gene expression at earlier stages. Our results argue that Chd4 regulates these aspects of retinal development via the NuRD (Nucleosome Remodeling and Deacetylase) complex (Xue et al., 1998; Zhang et al., 1998). Along with Class I CHD enzymes (Chd3/4/5), NuRD also includes class I histone deacetylases (Hdac1/2/6). Accordingly, Chd4-dependent shifts in rod versus Müller production were recapitulated by culturing retinal explants in the presence of pharmacological Hdac inhibitors, or by misexpressing a construct designed to interfere with NuRD assembly (Mattar et al., 2021).

While Chd4 is the key nucleosome remodeller within the NuRD complex in neural progenitors (Nitarska et al., 2016; Shah et al., 2025), Chd4 has more recently been associated with a second chromatin remodelling complex called ChAHP. (Ostapcuk et al., 2018). ChAHP consists of the ATP-dependent nucleosome remodeller <u>CH</u>D4, the zinc finger and homeodomain transcription factor <u>A</u>DNP, and the <u>HP</u>1 (Cbx1/3/5) heterochromatic proteins. ChAHP subunits are prominently associated with neurodevelopmental disorders (NDDs). In humans, *ADNP* is among the most frequently *de novo* mutated genes associated with autism spectrum disorder (ASD) and is also linked to intellectual disability (ID) (D’Incal et al., 2023; De Rubeis et al., 2014; Deciphering Developmental Disorders, 2017; Helsmoortel et al., 2014; Satterstrom et al., 2020; Stessman et al., 2017). Helsmoortel-Van der Aa syndrome (HVDAS; also known as ADNP syndrome; OMIM: 615873), is typically driven by *de novo* heterozygous truncating mutations (Van Dijck et al., 2019). Similarly, *de novo* mutations in *CHD4* are linked to Sifrim-Hitz-Weiss syndrome (SIHIWES; OMIM: 617159), which again features ID and more rarely ASD (Karimi et al., 2025; Sifrim et al., 2016; Weiss et al., 2020; Weiss et al., 2016). SIHIWES and HVDAS therefore share some common phenotypic features, and they exhibit a very similar episignature in blood DNA methylation profiles (Karimi et al., 2025).

Phenotypic data also hint that ChAHP may be important for visual system development. For example, SIHIWES is associated with optic nerve abnormalities (Karimi et al., 2025), and defects in visual functions have also been reported in HVDAS (Harutyunyan et al., 2026; Levy et al., 2026) - as well as in heterozygous *Adnp* mutant mice (Karmon et al., 2022). Moreover, an HVDAS case study reported mild foveal hypoplasia in one individual (Gale et al., 2018), suggesting a potential link between *ADNP* and retinal development. However, the *requirement* for ChAHP in neurodevelopment has been difficult to study. In pluripotent cells, *Adnp* mutants could not be differentiated into neural progenitors (Ostapcuk et al., 2018; Sun et al., 2020b; Yan et al., 2022), and *Chd4* mutation rapidly led to cell death (Stevens et al., 2017). Similarly, in mice, germline *Adnp* mutants died at approximately embryonic day (E) 9, and exhibited a failure to generate anterior neural tissue (Pinhasov et al., 2003), while *Chd4* mutation is lethal at pre-implantation stages (O’Shaughnessy-Kirwan et al., 2015). Brain-wide *Chd4* conditional knockouts (cKOs) were generated using the *Nestin-Cre* driver, but died at perinatal stages, precluding later analyses (Nitarska et al., 2016). The severity and heterogeneity of these phenotypes therefore led us to generate novel conditional mutants. Examining the telencephalon, we previously generated *Adnp* and *Chd4* cKOs using the *Emx1-Cre* driver. We found that both cKOs survived and exhibited highly similar neocortical phenotypes (Clémot-Dupont et al., 2024; Larrigan et al., 2026; Larrigan et al., 2023), arguing that Chd4 might contribute to neural development primarily through the ChAHP complex. However, it remains unclear whether this might be generally true in other regions of the nervous system – including the developing retina.

Here, we used conditional genetic models to study how ChAHP regulates retinal development. We confirm that the ChAHP complex is present in the developing retina, with Adnp and Chd4 proteins co-expressed ubiquitously in all cell types. Co-immunoprecipitations readily revealed an interaction between the two proteins. However, while *Adnp* and *Chd4* cKO phenotypes were nearly identical in the developing neocortex (Clemot-Dupont et al., 2025), they differed in the retina. We found that *Adnp* cKOs were hypoplastic but did not exhibit the marked shifts in cell fate seen in the *Chd4* cKO retina. To better understand this phenotypic divergence, we examined cut&run-seq datasets, and observed extensive co-recruitment of Adnp and Chd4 to Ctcf-occupied loci. Next, we performed scRNA-seq experiments to examine gene misregulation. Both *Adnp* and *Chd4* cKO cells exhibited a similar shift in gene expression, and co-clustered together. Some genes appeared to be coordinately misregulated in both mutants. However, *Chd4* cKOs exhibited much more extensive distortions in comparison to *Adnp* cKOs, suggesting that NuRD has a greater importance in retinal gene regulation. Together, these data reveal a connection between ChAHP and retinal hypoplasia, but suggest that retinal cell fate specification is largely ChAHP-independent.

## Results

### ChAHP complex expression during retinal development

To examine how ChAHP subunits are expressed in retinal development, we first consulted bulk and single-cell expression databases. *Adnp* transcript expression was not observed in many of these datasets. Looking more closely, we found that the *Adnp* exons are doubly annotated in the mouse genome, with a second readthrough transcript (NR_132310) consisting of the *Adnp* exons spliced to the transcript of an upstream gene called *Dpm1*, which encodes a biosynthetic enzyme. The readthrough transcript is classified as a lncRNA (Fig. S1). The presence of this second annotation in mm9, mm10, and mm11 genomes therefore creates a multi-mapping problem, leading to an apparent absence of expression in many RNA-seq datasets if/when ambiguously assigned reads are discarded.

We therefore utilized immunohistochemistry to examine the spatiotemporal expression profile of Adnp and Chd4 during retinal development using knockout-validated antibodies (Clemot-Dupont et al., 2025; Shah et al., 2025). We found that both proteins were ubiquitously expressed in all retinal cell types. At embryonic stage (E) 11, Adnp and Chd4 were colocalized within the nuclei of virtually every cell including Pax6+ RPCs (Fig. 1A-D). At E13.5 and 15.5, both Adnp and Chd4 levels remained ubiquitously co-expressed in dividing progenitors, but we observed elevated expression in postmitotic neurons - especially within the ganglion cell layer (GCL; Fig. 1E-H). At P1 and P5, we noted that Adnp expression was also elevated in mitotic cells at the apical surface of the developing retina (Fig. 1I-N) and also remained expressed within postmitotic Ki67-negative cells within the outer neuroblastic layer (ONBL; Fig. 1M-P). At later stages, the expression of Adnp and Chd4 in rod photoreceptors in the nascent outer nuclear layer (ONL) was weaker in comparison to cone photoreceptors (Fig. 1Q-V), and both proteins were maximally expressed within inner nuclear layer (INL) and GCL neurons (Fig. 1Q-V).

**Fig. 1.**
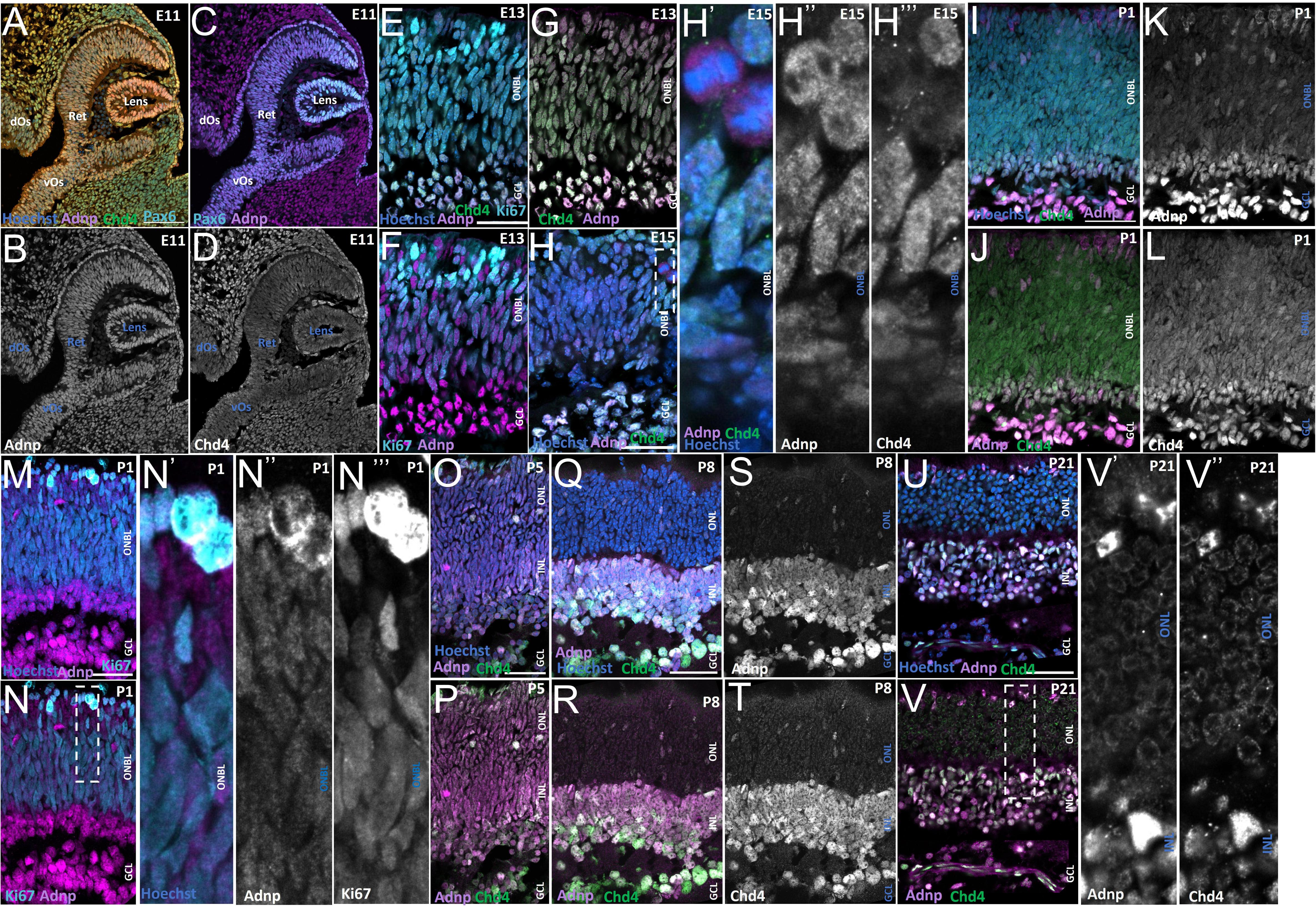
Adnp and Chd4 are co-expressed during retinal development. Retinal cryosections were stained with anti-Adnp (Magenta), anti-Chd4 (green), anti-Pax6 (Cyan), anti-Ki67 (Cyan) and the nucleus was counterstained by Hoechst (Blue). (A-H) Adnp and Chd4 co-expression at E11 (A-D), E13 (E-G), and E15 (H). (I-N) Adnp and Chd4 in proliferating progenitors at P1. (O-T) Adnp and Chd4 expression at later postnatal stages. (U, V) Adnp and Chd4 expression in the fully developed P21 retina. Dashed boxes indicate zoomed regions in (H, N, V). Ret, retina; Le, lens; Di, diencephalon; dOs, dorsal optic stalk; vOs, ventral optic stalk; ONBL, outer neuroblastic layer; ONL, outer nuclear layer; INL, inner nuclear layer; GCL, ganglion cell layer. Scale bars: 50 μm.

Since Adnp and Chd4 were extensively co-localized, we next performed co-immunoprecipitation (coIP) and mass spectrometry (MS) analysis to determine whether Adnp and Chd4 proteins interact *in vivo*. We used the same knockout-validated antibodies to co-IP Adnp and Chd4 from P0 retinal tissue extracts. In Adnp coIPs, Chd4 was the most predominant interacting partner, and we also observed significant enrichment of the known ChAHP subunit CBX3 (Fig. 2A). Conversely, Adnp was among the most abundant proteins identified within the Chd4 interactome (Fig. 2B). However, as expected, the CHD4 interactome additionally contained many subunits of the NuRD complex that were absent from ADNP coIPs, such as GATAD2A/B and HDAC1/2, MBD3, RBBP4/7, and MTA1/2/3, demonstrating the fidelity of the interactomes (Fig. 2B; Supplemental data file 1). Reciprocal coIP/western blots of Adnp and Chd4 readily confirmed the presence of the ChAHP complex within the developing retina (Fig. 2C).

**Fig. 2.**
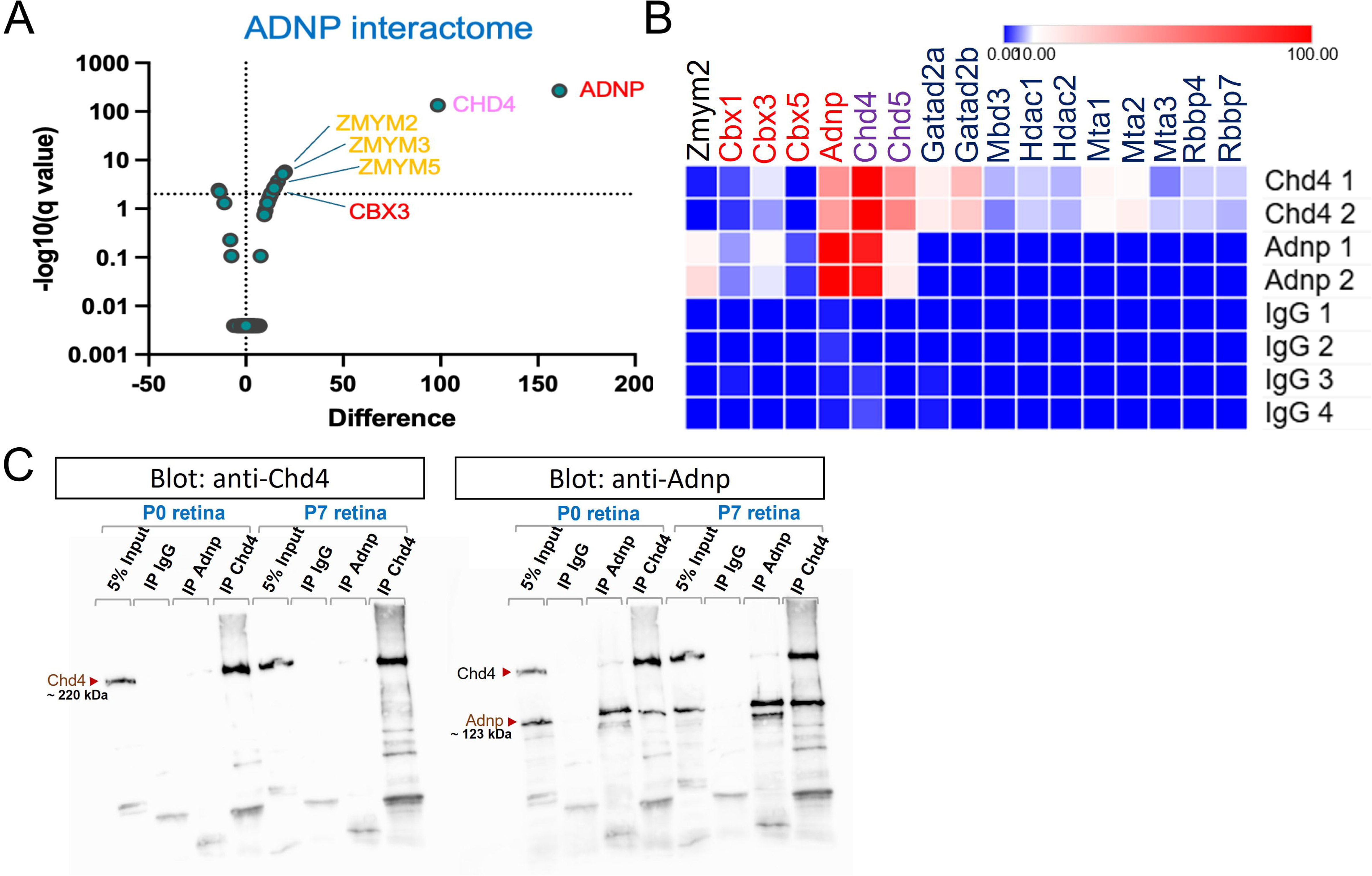
Adnp and Chd4 interact to form the ChAHP complex in the mouse retina. (A) Volcano plot of Adnp coIP/MS results. Chd4 was the most predominant interacting partner in the Adnp interactome. (B) Heatmap of selected ChAHP and NuRD subunits in coIP/MS experiments. Color intensity represents the peptide counts identified in MS, ranging from low/no interaction (blue) to strong interaction (red) as indicated. (C) Adnp and Chd4 IP/westerns from P0 or P7 retinas. The membrane was first probed with anti-Chd4 antibodies and subsequently reprobed with anti-Adnp antibodies without stripping the membrane.

### Conditional ablation of ChAHP subunits in the developing retina

Previous studies have reported that germline *Adnp* knockout mice are non-viable (Pinhasov et al., 2003). We therefore generated a conditional *Adnp* allele using CRISPR/Cas9 to introduce the loxP sites on either side of the exon 5 of *Adnp* (Clemot-Dupont et al., 2025), which contains all of Adnp’s DNA binding domains (Fig. 3A). To test the importance of the ChAHP complex in retinal development, we used *Chx10-Cre* (Rowan and Cepko, 2004) to inactivate *Adnp* specifically in the neural retina beginning at E11, generating wild-type (WT; *Adnp^+/+^* and/or Cre-negative), *Adnp* conditional heterozygote (cHET; *Adnp^Flox/+^*; *Chx10-Cre+*), or *Adnp* cKO (cKO; *Adnp^Flox/Flox^*; *Chx10-Cre+*) mice. To validate the *Adnp* cKO model, we performed western blotting analysis using P0 retinal tissue from the three genotypes. As expected, Adnp protein was strongly reduced in *Adnp* cKO samples relative to the other genotypes (Fig. 3B-C). Interestingly, immunohistochemistry on P1 retinas confirmed the ablation of Adnp protein, but also revealed ∼50% reduction in Chd4 protein levels. Examining *Chd4* cKOs, we observed a corresponding reduction in Adnp protein levels (Fig. S2). These data suggest that the integrity of the ChAHP complex might be important for subunit levels and/or stability.

**Fig. 3.**
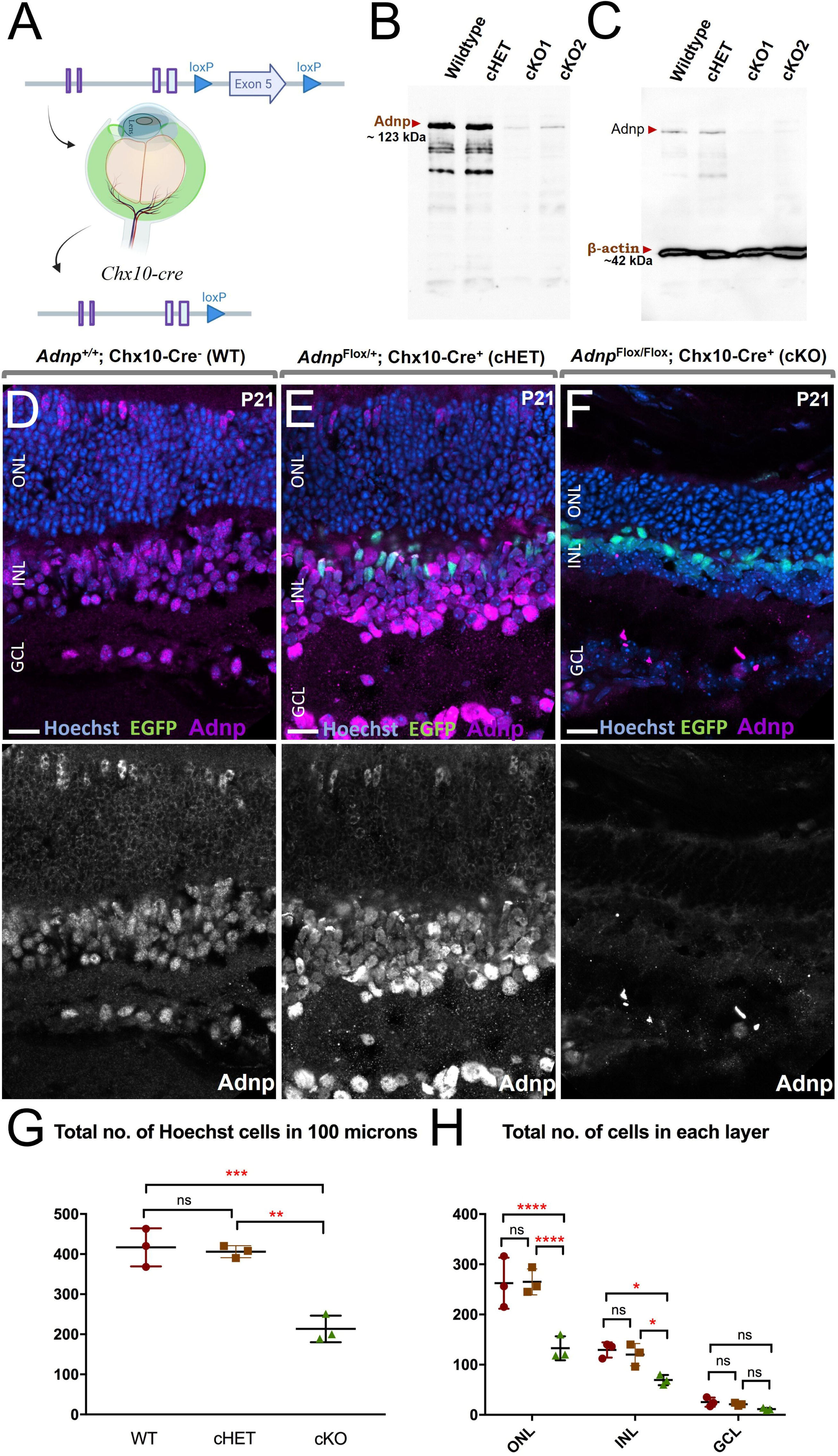
Conditional deletion of *Adnp* during retinal development leads to hypocellularity. (A) The *Chx10-Cre* driver directs the inactivation of Adnp, specifically in RPCs beginning at E10.5. (B) Western analysis of Adnp protein expression and (C) β-actin as a loading control in wild-type, cHET and cKO retinas. (D-F) Adnp immunostainings on P21 WT (D), cHET (E) and cKO retinas (F). Scale bar: 20μm. (G, H) Total cell counts (G), as well as cell counts by retinal layer (H). *p<0.05, **p < 0.01, ***p < 0.001, \*\*\*\**p* < 0.0001 via one-way ANOVA with the Tukey/Kramer post-hoc test.

### Adnp is required for retinal growth

We next examined *Adnp* cKOs at P21. As expected, Adnp protein expression was significantly lost in cKO retinas (Fig. 3D-F). We did not find any morphology changes in cHET retinas, but the cKO retinas were considerably smaller. Cell counts revealed clear reductions in the ONL and INL (Fig. 3G-H). However, the proportion of cells in each layer was not significantly altered (Fig. S3).

To determine how Adnp loss affects retinal cell-type composition, we used marker proteins to count each cell-type, including cones and rods (distinguished via cone arrestin; CA; Fig. 4A, G, H), horizontals (calbindin at the outer plexiform layer; Fig. 4B, I), bipolars (Otx2 in the INL; Fig. 4C, J), Müller glia (radially polarized Sox2+ nuclei in the central INL; Fig. 4D, K), amacrines and ganglion cells (Pax6; Fig. 4E, L; Rbpms Fig. 4F; M). The data revealed significant reductions in many cell types, while offsetting increases in specific cell types were not observed, suggesting that Adnp does not alter cell type specification. Accordingly, when data were reanalyzed to examine the overall proportions of each cell type, no change in proportionality was observed (Fig. S3). Interestingly, we also observed small numbers of mismigrated cells in the *Adnp* cKO, including Müller glia and bipolars (Fig. 4C-D, Fig. S4). Strikingly, a few ectopic ganglion cells were found in ONL and INL (Fig. 4F).

**Fig. 4.**
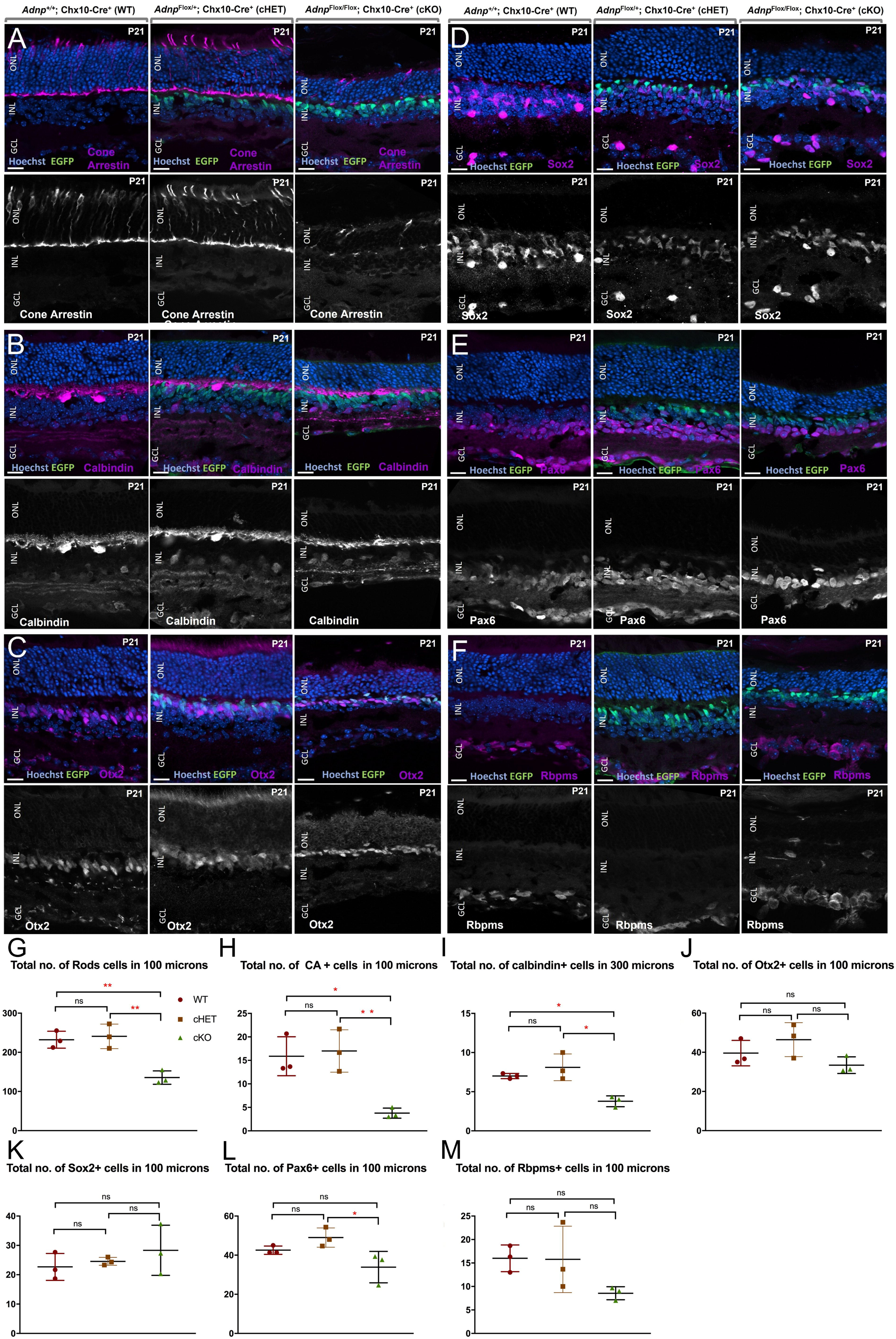
Loss of Adnp leads to comparable reductions in all cell types. (A-F) Cell-type marker expression in WT, *Adnp* cHET and *Adnp* cKO retinas at P21. Cone Arrestin (A; cones), Calbindin (B; horizontals, amacrines, RGCs), Otx2 (C; bipolar cells), Sox2 (D; Müller glia and cholinergic amacrines), Pax6 (E; amacrines, RGCs), Rbpms (F; RGCs). ONL, outer nuclear layer; INL, inner nuclear layer: GCL, ganglion cell layer. Scale bars: 20 μm. (G-M) Absolute counts for retinal cell types defined by the above markers. See also the text for further explanation. *p<0.05, **p < 0.01, via one-way ANOVA with the Tukey/Kramer post-hoc test.

The hypocellularity exhibited by the *Adnp* cKO contrasts with the marked shifts in cell type composition and defective lamination that we previously observed in the *Chd4* cKO (Shah et al., 2025). We therefore examined RPC proliferation and cell death in *Adnp* versus *Chd4* cKOs by performing Ki67 and activated-caspase3 immunostainings on P1 retinas (Fig. 5A-C). We found that the *Adnp* cKO exhibited significant reductions in total cell numbers, and in the absolute numbers of Ki67-positive progenitors (Fig. 5I, J). By comparison, these metrics remained unchanged in *Chd4* cKOs versus wildtypes (Fig.5 D-J) as previously reported (Shah et al., 2025). Notably, there was a significant reduction in the number of mitotic cells (strong apical Ki67+ cells) in the *Adnp* cKOs compared to *Chd4* cKOs (Fig. 5C, J). We also found significant increases in activated caspase-3-positive cells in both cKOs (Figure 5E-G, K). These findings suggest that ChAHP is necessary for the survival of RPCs.

**Fig. 5.**
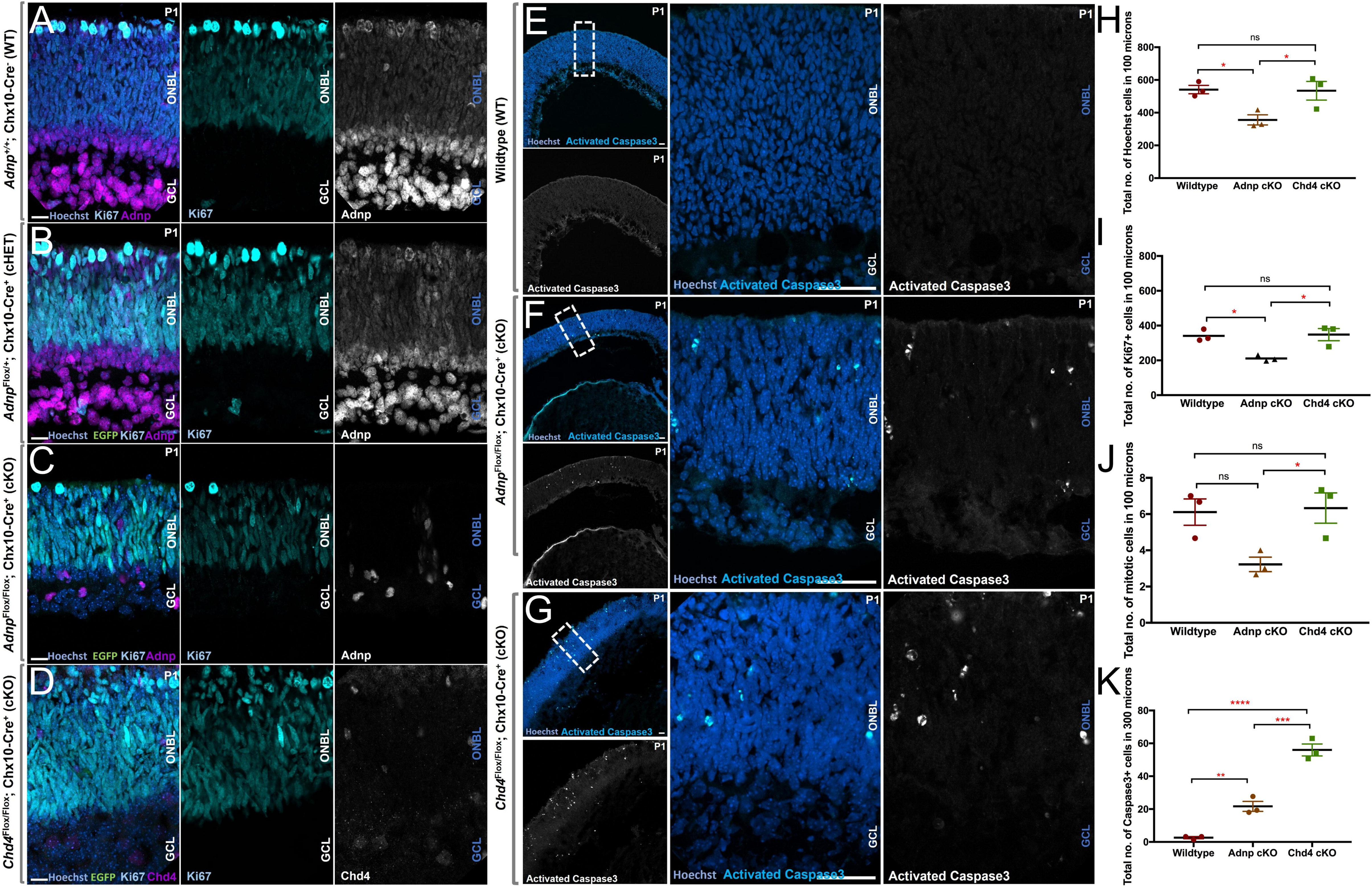
Loss of Adnp leads to hypoproliferation and increased cell death. (A-D) Retinal cryosections of P1 WT (A), *Adnp* cHET (B), *Adnp* cKO (C) and *Chd4* cKO (D) retinas stained with anti-Ki67, anti-Adnp or anti-Chd4, and co-stained with Hoechst. Scale bar: 20μm. (E-G) Retinal cryosections of P1 WT (E), *Adnp* cKO (F) and *Chd4* cKO (G) stained with anti-activated caspase3, and co-stained with Hoechst. Dashed boxes indicate zoomed insets. Scale bar-50 μm for lower magnification images and 20 μm in zoomed images. (H-K) Cell counts of Hoechst+ cells (H), Ki67 positive cells (I), mitotic cells (J) and activated caspase-3 -positive cells (K) in WT, *Adnp* cKO and *Chd4* cKO retinas. *p<0.05, **p < 0.01, ***p < 0.001, \*\*\*\**p* < 0.0001 via one-way ANOVA with the Tukey/Kramer post-hoc test.

### Coordinate and divergent effects of Adnp and Chd4 on gene regulation

The ChAHP complex has been well studied in the context of pluripotent cells, where it has been shown to occupy heterochromatic regions of the genome that contain Ctcf motifs (Kaaij et al., 2019; Ostapcuk et al., 2018; Yan et al., 2022). By contrast, in more differentiated contexts such as the developing cortex or in T cells, Adnp has been shown to bind to euchromatic elements such as promoter and enhancer elements (Ferreira et al., 2023; Markenscoff-Papadimitriou et al., 2021; Wulfridge et al., 2026). Moreover, Adnp has been previously suggested to associate with a variety of non-ChAHP interaction partners, with the BAF nucleosome remodelling complex being particularly relevant to gene regulation (D’Incal et al., 2023). Thus, ChAHP –dependent and independent functions might be expected – both for Adnp and Chd4, with the latter prominently linked to the NuRD complex.

Since ChAHP functions are poorly understood in the developing nervous system, we decided to compare *Adnp* and *Chd4* cKOs at the transcriptomic level. We used multi-seq to profile 3 wild-type (cre-negative) and 3 *Adnp* cKO retinas at P1. Multiseq uses barcoded oligonucleotides to mark individual replicates, which can then be mixed and run together in a single 10X Genomics experiment. The barcodes include runs of A-residues, allowing them to be captured by the oligo-dT primers arrayed on the beads. Next, we integrated the Adnp dataset with a previously generated multiseq dataset that similarly consists of 3 control vs. 3 *Chd4* cKO P1 retinas. We performed demultiplexing, filtered low quality and contaminant cells, and performed batch correction, leaving 9,580 cells from the prior Chd4 dataset, along with 4,521 cells from the new Adnp experiment (Table S1).

Next, cell types were clustered using the Leiden algorithm, and we manually annotated each cluster based on marker gene expression (Fig. S5). UMAP projection revealed a relatively simple structure, with a group of primary progenitor cells connected to bifurcating photoreceptor cell or amacrine cell trajectories in a ‘Y-like’ configuration, similar to what we previously observed with the Chd4 dataset alone (Shah et al., 2025). We noted that cells from both experiments and all 12 individual replicates generally populated these clusters in a similar fashion (Fig. 6B, C; Fig. S6). As a quality check, we also confirmed that *Adnp* transcript was reduced specifically in the *Adnp* cKO replicates (Fig. 6D), while *Chd4* was reduced in *Chd4* cKOs, albeit to a much lesser degree (Fig 6E) – consistent with the fact that the conditional *Chd4* allele leaves the 3’ exons intact (Williams et al., 2004). Similarly to our previous findings (Clemot-Dupont et al., 2025; Shah et al., 2025), we observed that *Chd3* and *Chd5* paralogs upregulated to compensate for *Chd4* ablation, whereas *Adnp2* was little expressed, and did not compensate for the loss of *Adnp* (Fig. 6E).

**Fig. 6.**
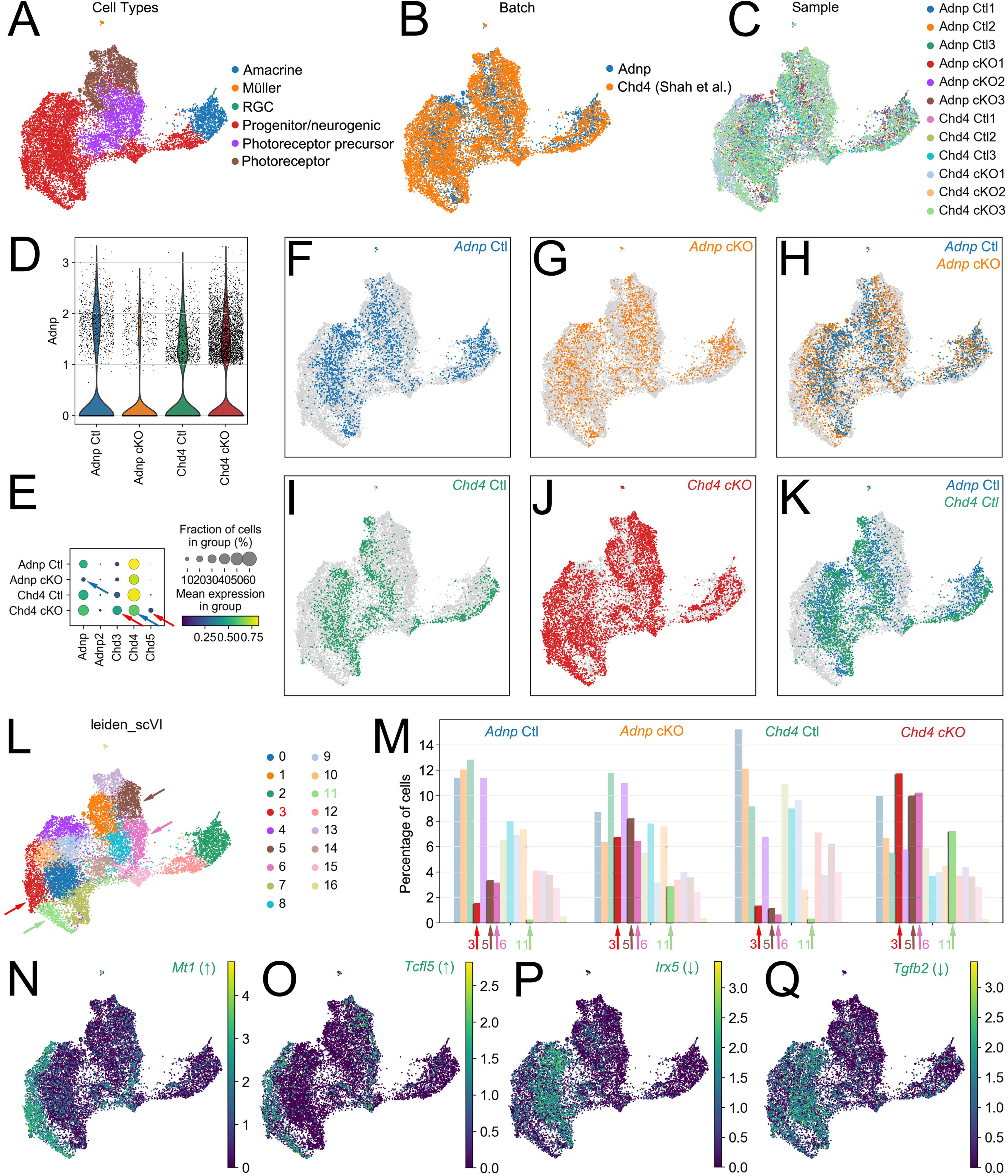
Coordinate transcriptomic shifts in *Adnp* and *Chd4* cKOs. (A-C) UMAP embedding of the combined scRNA-seq dataset color-coded for cell-type (A), experimental batch (B), or replicate (C). (D, E) Expression of *Adnp*, *Chd4*, and relevant paralog transcripts in *Adnp* cKOs, *Chd4* cKOs, or controls. (F-K) UMAP embedding and overlap of *Adnp* cKOs, *Chd4* cKOs, or controls as indicated. (L, M) Shifts in cell distributions among Leiden clusters in *Adnp* cKOs, *Chd4* cKOs, or controls as indicated. Arrows indicate clusters 3, 5, 6, and 11, which are enriched in cKO cells. (N-Q) UMAP embeddings of DEGs, including transcripts upregulated in cKO cells (*Mt1*, N; *Tcfl5*; O), or downregulated in cKO cells (*Irx5*, P; *Tgfß2*, Q).

Examining individual genotypes, we noted that both *Adnp* and *Chd4* cKO cells occupied a group of clusters that were relatively devoid of control cells (Fig. 6F-K). These clusters lay adjacent to progenitor clusters as well as to photoreceptor precursors. The cKO –specific clusters corresponded to Leiden clusters 3 and 11 (progenitors) and 5 and 6 (photoreceptor precursors). The proportions of cells in these clusters jumped noticeably in both *Adnp* and *Chd4* cKOs (Fig. 6L, M), suggesting coordinate shifts in gene expression. We next examined the *Adnp* and *Chd4* cKOs to identify groups of differentially expressed genes (DEGs). Examining some DEGs individually, we found that differentially upregulated genes such as *Mt1* and *Tcfl5* marked the novel cKO clusters specifically (Fig. 6N, O). Moreover, downregulated progenitor DEGs such as *Irx5* and *Tgfb2* were markedly reduced in the novel clusters relative to the parallel progenitor clusters occupied by control cells (Fig. 6P, Q).

Next, we examined DEGs systematically. In pseudobulk analyses of the *Chd4* cKO replicates, we observed 540 DEGs, similar to the results obtained for *Chd4* cKOs prior to merging datasets (Shah et al., 2025). By contrast *Adnp* cKOs exhibited only 42 DEGs, the majority of which were also differential in the *Chd4* cKO (Fig. 7A; Supplemental data file 2). Correlation between the two cKOs appeared relatively weak, with many DEGs from the *Adnp* cKO exhibiting relatively little alteration in the *Chd4* cKO (Fig. 7B). Thus, while analysis of scRNA-seq clusters suggested that *Adnp* and *Chd4* cKOs exhibited transcriptomic resemblance, only a few coordinately regulated DEGs were identified. To gain insight into how ChAHP regulates the genome, we next performed cut&run-seq on P0 wild-type retinas using the same knockout-validated Adnp antibody. Again, we compared these results versus our published cut&run-seq for Chd4 (Shah et al., 2025). In accordance with previous findings, Adnp peaks were found to be over-represented with GC rich motifs that bore some resemblance to a Ctcf half-site (Fig. 7C). We therefore performed cut&run-seq for Ctcf. Comparing peaksets, we noticed considerable overlap between Adnp, Chd4, and Ctcf. ∼40% of the Adnp peaks were co-occupied by both Ctcf and Chd4, with another ∼40% co-occupied by Ctcf alone (Fig. 7D). Benchmarking revealed strong correspondence between our Ctcf cut&run-seq data and previously published Ctcf ChIP-seq data (Aldiri et al., 2017) (Fig. S7).

**Fig. 7.**
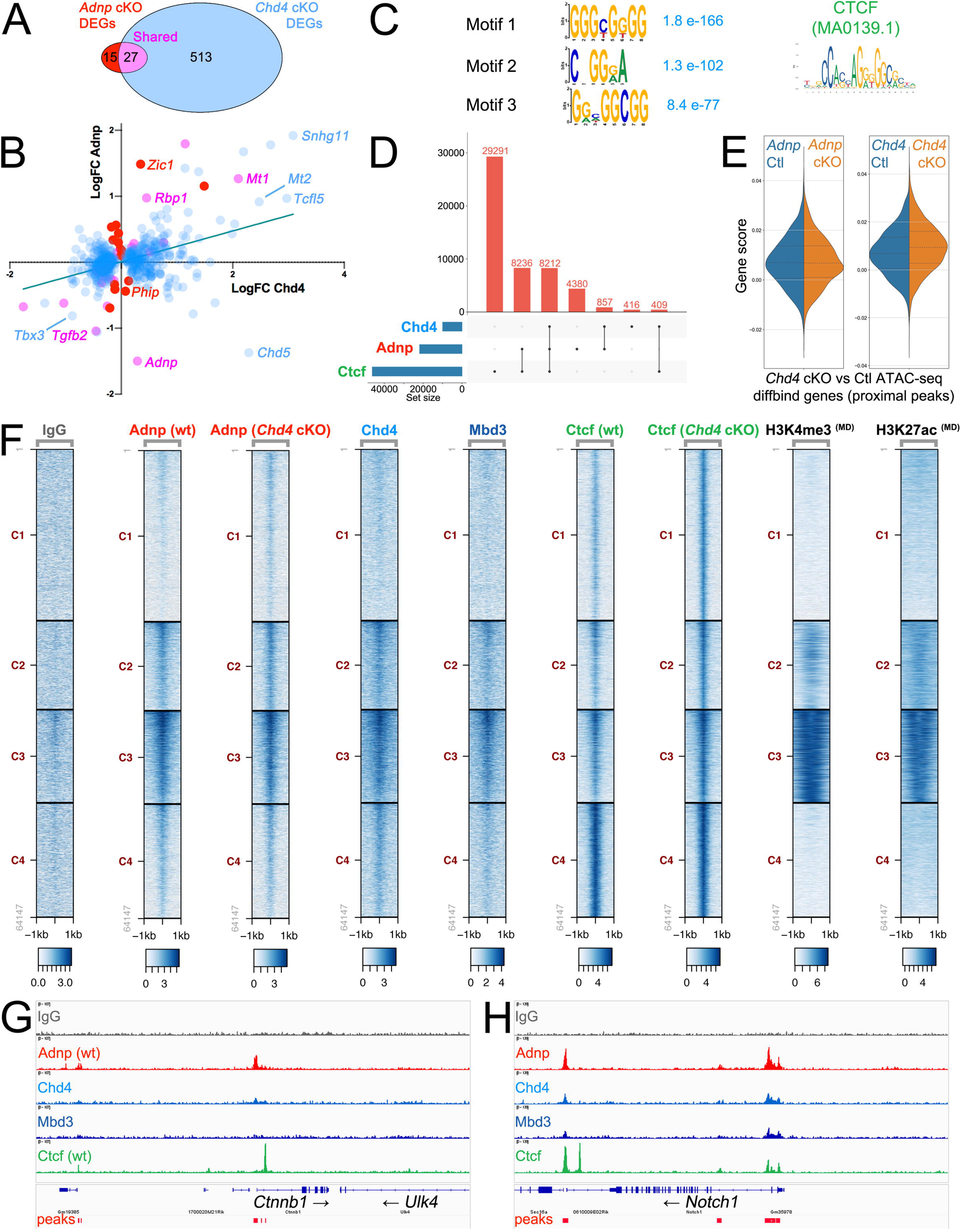
Retinal gene regulation by Chd4 and Adnp. (A) Venn diagram comparing DEGs identified in *Adnp* or *Chd4* cKOs versus their respective controls. (B) Scatterplot demonstrating the coordinate change in transcript expression in *Adnp* or *Chd4* cKOs versus their respective controls (red: *Adnp* DEGs; blue *Chd4* DEGs; purple: shared DEGs). (C) Comparison of top 3 overrepresented motifs present in Adnp cut&run-seq called peaks versus a canonical Ctcf motif. (D) Comparison of Adnp, Chd4, and Ctcf peak numbers from retinal cut&run-seq. (E) Gene score for genes previously found to be associated with differential accessibility in *Chd4* cKO retinas (Shah et al., 2025). (F) Cut&run-seq datasets derived from P0 retinas for the indicated proteins/marks. MD: Michael Dyer laboratory (Aldiri et al., 2017). (G, H) Example of Adnp, Chd4, and Mbd3 (NuRD subunit) recruitment to the *Ctnnb1* or *Notch1* loci.

To gain further insight into how Adnp regulates Chd4 targets, we next examined genes that we previously identified to be associated with differential accessibility in *Chd4* cKOs (Shah et al., 2025), suggesting that they may be direct targets of Chd4-dependent nucleosome remodelling. We generated a gene scoring module for this group of targets. Interestingly, while there was an obvious upregulation of this module in *Chd4* cKOs, no obvious shift in the expression of these genes was evident in the *Adnp* cKO (Fig. 7E).

Taken together, phenotypic and transcriptomic data suggest that Chd4 has a much stronger effect on retinal gene regulation versus Adnp. One interpretation is that Chd4 might regulate genes independently of Adnp/ChAHP - acting via the NuRD complex. To examine this in more detail, we compared ChAHP (Adnp, Chd4) occupancy with NuRD (Chd4, Mbd3), and Ctcf, which revealed extensive co-occupancy on the genome – particularly at promoters marked by H3K27ac and H3K4me3 (Fig. 7F). We additionally examined published data for the BAF subunit Brg1 (Aldiri et al., 2015), but observed no overlap between Brg1 and Adnp occupancy (Fig. S8), in accordance with the lack of protein interactions observed in coIP/Ms experiments. Interestingly, in *Chd4* cKOs, we observed increased Ctcf occupancy relative to wild-type, which was reflected in an increase in called peaks from ∼45,000 up to ∼64,000. This result matches the increase in Ctcf motif footprints in the *Chd4* cKO that we previously observed in ATAC-seq data (Shah et al., 2025), as well as published results from cerebellar granule neurons (Goodman et al., 2020). Accordingly, inspection of occupied loci revealed that Adnp-occupied genes such as *Ctnnb1* and *Notch1* tended to exhibit correlating signal for Chd4, Mbd3, and Ctcf (Fig. 7G, H). Taken together, these data suggest a surprising correspondence between NuRD and ChAHP occupancy, and hint that Chd4 remodelling activity may oppose the retention of Adnp on the genome.

## Discussion

The importance of the ChAHP complex in neurodevelopment is underscored by genetic associations between its constituent subunits and neurodevelopmental disorders. *ADNP* is of particular importance in this regard - ranking among the most commonly *de novo* mutated genes associated with autism spectrum disorder (Satterstrom et al., 2020). Despite this fact, it remains unclear how ChAHP regulates nervous system development. This is partly due to the fact that the complex was first described relatively recently (Ostapcuk et al., 2018), and in part because genetic knockouts arrest early in development (O’Shaughnessy-Kirwan et al., 2015; Pinhasov et al., 2003). Even in pluripotent stem cell models, *ADNP* mutation precludes neural differentiation (Ostapcuk et al., 2018; Sun et al., 2020b; Yan et al., 2022), necessitating conditional genetic strategies.

In humans, initial characterization of neurodevelopmental disorders associated with *ADNP* (HVDAS) and *CHD4* (SIHIWES) suggested some divergence in neurodevelopmental phenotypes. While both syndromes are associated with intellectual disability, *ADNP* is highly associated with autism spectrum disorder (Helsmoortel et al., 2014; Satterstrom et al., 2020). Moreover, SIHIWES features macrocephaly and microcephaly (Weiss et al., 2020; Weiss et al., 2016), which have not been linked to HVDAS. However, both syndromes share similar DNA methylation episignatures, and SIHIWES has more recently been associated with autism (Karimi et al., 2025).

Visual deficits are also linked to both syndromes. Optic nerve abnormalities were observed in some SIHIWES patients (Karimi et al., 2025). In HVDAS, sensory and visual abnormalities are common (Harutyunyan et al., 2026; Levy et al., 2026), although the mechanisms underlying these deficits remain unclear. One report identified mild foveal hypoplasia in a single HVDAS case (Gale et al., 2018), suggesting a potential role for *ADNP* in retinal development. However, visual systems deficits could conceivably originate in the brain - in line with our previous study that demonstrated roles for *Adnp* in cortical neurogenesis (Clemot-Dupont et al., 2025).

Here, we used conditional genetics to dissect the contributions of ChAHP subunits to retinal neurogenesis. In the *Adnp* cKO, we observed reductions in proliferation and elevations in apoptosis that likely underlie retinal undergrowth. A functional link between ChAHP subunits and elevated cell death has been well established by prior studies (Koulle et al., 2026; Nitarska et al., 2016; Ostapcuk et al., 2018; Sun et al., 2020a). More unexpectedly, *Adnp* cKOs exhibited decreased proliferation, while *Chd4* cKOs did not. This is the first phenotype found in *Adnp* cKOs that was not also found in *Chd4* cKOs. Adnp might therefore regulate RPC proliferation in a ChAHP -independent manner. However, as Chd3/5 were previously shown to be capable of forming ChAHP complexes with Adnp (Tabar et al., 2025), it might instead reflect genetic compensation. Chd3/5 are not ordinarily expressed in RPCs, but we previously showed that both subunits upregulate in *Chd4* cKOs (Shah et al., 2025). Additional work will be required to distinguish between these possibilities.

Similarities and differences were also observed in transcriptomic and genomic datasets. Correspondence between *Adnp* and *Chd4* cKOs was observed in scRNA-seq clustering, where cKO cells from both mutants were coordinately shifted into novel clusters. Moreover, most of the DEGs identified in *Adnp* cKOs were also differentially regulated in *Chd4* cKOs. However, *Chd4* cKOs exhibited much more extensive gene misregulation. Batch effects across the scRNA-seq experiments could potentially contribute to these findings, since the *Adnp* and *Chd4* cKOs were processed separately. However, genes found to be differentially accessible in the *Chd4* cKO based on ATAC-seq experiments exhibited no sign of misregulation in *Adnp* cKOs – irrespective of significance testing. Moreover, *Chd4* cKOs exhibited much stronger distortions at the phenotypic level in accordance with the much larger number of DEGs observed in these mutants. Since Chd4 also regulates the genome via the NuRD complex, a stronger phenotype was expected. However, we were nonetheless surprised by the degree to which *Adnp* and *Chd4* cKO retinal phenotypes diverge – especially since comparisons between *Adnp* and *Chd4* germline heterozygotes and neocortical cKOs have not revealed clear phenotypic differences (Clemot-Dupont et al., 2025) – even at the level of behavior (Larrigan et al., 2026; Larrigan et al., 2023). These findings suggest that the developing brain might be particularly dependent on ChAHP functions, whereas the retina appears to be much more sensitive to disruptions in NuRD-dependent chromatin remodelling. Taken together, our study dissects the role of the ChAHP complex on retinal development and gene regulation, and we think that our observations provide insight into the mechanisms that might undermine the visual system in HVDAS and SIHIWES syndromes.

## Materials and Methods

### Animals

Animal work was conducted according to the guidelines of the Canadian Council of Animal Care and the uOttawa Animal Care and Veterinary Service (ethical protocols OHRI-3949, OHRI-4029, and OHRI-4761). Generation of the *Adnp^Flox^* allele was previously described (Clemot-Dupont et al., 2025). *Chd4^Flox^* mice were generously donated by Dr. Katia Georgopoulos (Williams et al., 2004), and backcrossed onto the C57BL/6J background (Jackson Labs; RRID: IMSR_JAX:000664). *Chx10-Cre-GFP* transgenic mice (RRID:IMSR_JAX:005105) (Rowan and Cepko, 2004) were generously provided by Dr. Catherine Tsilfidis. Genotyping primers are presented in Table S2. Both males and females were used for all experiments.

### Histology

For embryonic stages, heads were collected and fixed in 4% PFA overnight at 4°C, followed by PBS washes, and then immersion in PBS/20% sucrose at 4°C overnight. After further washing, heads were immersed in a 1:1 solution of PBS/20% sucrose: OCT overnight at 4°C. Tissues were embedded and frozen in the same OCT/sucrose mix. For postnatal stages (P0 - P21), a small slit was introduced in the anterior chamber and the cornea was removed, leaving the lens intact. This was followed by 4% PFA fixation for 1-2 minutes. After fixation, tissues were washed in PBS, and transferred into PBS/20% sucrose at 4°C for 2-3 hours or overnight. Retinas were embedded in OCT and frozen.

12µm coronal cryosections were collected onto Superfrost Plus slides (Fisher) and processed for immunofluorescence as described previously (Shah et al., 2025). Briefly, tissue sections were washed in PBS followed by antibody incubation at 4°C overnight. in blocking buffer (PBS supplemented with 0.4% Triton X-100, 3% w/v BSA, and Hoechst 33342). After washing with PBS, secondary antibodies were incubated in blocking buffer for one hour at room temperature. See Table S3 for antibodies and dilutions. Coverslips were mounted using the Mowiol mounting media (12% w/v Mowiol 4-88, 30% w/v glycerol, 120mM Tris-Cl pH 8.5, 2.5% DABCO).

### Imaging and Cell counting

Images were acquired on an LSM900 confocal microscope (Zeiss). *Chx10-Cre-GFP* mosaicism was monitored by inspecting GFP epifluorescence during developmental stages. For stages beyond P8, we monitored mosaicism by co-staining sections with anti-Adnp antibodies. We avoided retinal fields that exhibited significant *Chx10-Cre-GFP* mosaicism. For each biological replicate, 3 optical sections (single Z planes) of the peripheral retina were used to generate cell counts. Images were further processed using Fiji (ImageJ).

### Statistical analysis

Statistical analysis was performed using GraphPad Prism version 8 (GraphPad) software. n-values and datapoints in graphs refer to biological replicates (animals). Statistical analyses were performed via one-way ANOVA with Tukey’s multiple comparison test or two-tailed unpaired t-tests as appropriate. We did not perform statistical analyses to predetermine sample sizes. Graphs depict mean ± standard error of mean (SEM).

### Co-immunoprecipitation

Retinal tissue was freshly microdissected and sonicated in RIPA buffer (10 mM Tris pH 8.0; 140 mM NaCl; 1mM EDTA; 0.5 mM EGTA; 1% Triton X-100; 0.1 % deoxycholate; 0.1 % SDS) supplemented with cOmplete™ EDTA-free Protease Inhibitors (11836170001; Roche). 30 µl of protein A or G paramagnetic beads (73778S or 9006S; New England Biolabs) were washed and then co-incubated with primary antibodies or naïve rabbit IgG and incubated at 4°C with rotation. Antibody information is presented in Table S3. Beads were washed and 2 mg of retinal protein lysate was added. Lysates were diluted to 1 ml with RIPA buffer with protease inhibitors and rotated overnight at 4°C. Next, the lysates were removed, and beads were transferred into fresh tubes. Beads were then washed 3x 30 min with RIPA buffer. For western analysis, proteins were eluted by boiling the paramagnetic beads in Laemmli Sample buffer (1610747; Bio-Rad) supplemented with 2-mercaptoethanol.

### Western Blot

Western blotting was performed as previously described (Mattar et al., 2021) with some modifications. P0 dissected retinas were homogenized in RIPA buffer with protease inhibitors (cOmplete, Mini, EDTA-free; 11836170001; Millipore Sigma) and incubated on ice for 15 minutes. Samples were carefully sonicated on ice and insoluble material was pelleted and discarded. The supernatant was collected, and the protein concentration was quantified using BCA assay. Approximately 40µg of protein lysate was separated on a 6-10% gradient SDS-PAGE gels and semi-dry transferred onto PVDF membranes (Millipore). Antibody incubations were performed using condensed milk powder as a blocking reagent. Chemiluminescence reactions were visualized using a Chemidoc Touch™ instrument (Bio-Rad). Antibodies and dilutions are presented in Table S3.

### LC–MS/MS analysis

CoIP experiments were performed as described above. MS was performed via on-bead tryptic digestion by the *Ottawa Hospital Research Institute Proteomics Core Facility* (Ottawa, Canada). Peptide samples were concentrated using a Vacufuge (Eppendorf) and with ZipTips (Millipore Sigma). LC-MS/MS was performed using a Dionex Ultimate 3000 RLSC nano HPLC (Thermo Scientific) and Orbitrap Fusion Lumos mass spectrometer (Thermo Scientific). Peptide mass/spectral fingerprinting was performed using MASCOT software version 2.7.0 (Matrix Science, UK). Observed spectra were matched against mouse sequences from SwissProt (version 2021-02), and against an in-house-curated database of common contaminants. The results were exported to Scaffold 5.0.1 (Proteome Software, USA). Identification of significantly enriched proteins was performed using Graphpad Prism 8 via the Holm-Sidak method, with alpha = 0.05.

### Cut&run-seq

Cut&run-seq was performed as previously described (Clemot-Dupont et al., 2025). Briefly, pairs of retinas from individual pups were dissected. Some samples were banked by slowly freezing them in tissue culture medium supplemented with 10% DMSO and stored at -80°C. Retinas were next mechanically dissociated by trituration with a pipette tip so as to maintain the integrity of surface glycoproteins necessary for adhesion to the concavanalin beads. Cut&run-seq was performed using the Cutana ChIC/CUT&RUN kit (14-1048; Epicypher), following the manufacturer’s instructions.

### Retinal dissociation

Retinas were dissociated using papain (0.003N NaOH, 100U papain solution (Worthington), 0.4% DNaseI (Millipore Sigma 04716728001), 2mg L-cysteine crystal in PBS) for 8 minutes at 37°C. Subsequently, papain buffer was aspirated and replaced with LO-OVO solution (LO-OVO (Bio Basic), 0.4% DNaseI in PBS). Thereafter, the retinal tissue was triturated and centrifuged for 11 minutes at 200g at room temperature, and cells were re-suspended in PBS.

### Multi-seq

Multi-seq was performed as described previously (Shah et al., 2025) with some modifications. P1 retinas from an *Adnp*/*Chx10-Cre* litter were dissociated. Approximately 250,000 dissociated cells per replicate were then barcoded by incubating with ‘anchor’ and ‘co-anchor’ lipid-modified oligonucleotides graciously provided by the Gartner lab. Barcode oligonucleotides were purchased from Integrated DNA Technologies (see Table S2). Individual replicates were co-incubated with barcode oligonucleotides and pooled into a single tube at a 1:1 ratio. Approximately 20,000 pooled cells were used in a single Chromium™ run (3′ Library & Gel Bead Kit v2, PN-120237, 10X Genomics). The resulting expression library FASTQs were processed using CellRanger (10X Genomics). The deMULTIplex workflow was used to perform quality control to remove doublets along with cells that lacked the barcodes.

Datasets were processed and integrated using Scanpy (Wolf et al., 2018) and single-cell variational inference modelling (scVI) (Lopez et al., 2018). Specifically, quality control was performed using Scanpy, and gene names beginning with mt-were defined as mitochondrial. Per-cell quality metrics, including the number of detected genes, total counts, and percentage of mitochondrial transcripts, were calculated using scanpy.pp.calculate_qc_metrics. Contaminant cells were identified and removed based on the expression of non-retinal genes. Outliers were filtered to retain cells with >500 detected genes, 1,000–15,000 total counts, and less than 5% mitochondrial transcripts using sc.pp.filter_cells. Following quality control, the 2 Multiseq datasets were concatenated for integration. To account for batch-associated variation, an scVI model was configured using raw gene counts as input, with each sequencing batch included as a categorical covariate. Continuous covariates included the percentage of mitochondrial and ribosomal transcripts, total counts, number of detected genes, a background gene expression score, and a sex-associated gene expression score. The background gene expression score was calculated for each cell by summing expression across the top 1,000 genes ranked by differential expression between the *Adnp* control and *Chd4* control groups. The sex-associated gene score was calculated by summing expression across a predefined panel of sex-associated genes, including *Xist*, *Tsix*, *Ddx3y*, *Eif2s3y*, *Kdm5d*, and *Uty*. The trained scVI model was then used to generate a batch-corrected latent representation for downstream analyses. Batch-corrected normalized counts were obtained using model.get_normalized_expression. Uniform Manifold Approximation and Projection (UMAP) (McInnes et al., 2018) was used to visualize the data in a reduced dimensional space. Cell clustering was performed using the Leiden algorithm (Traag et al., 2019). Differential expression between groups was performed using a Wilcoxon rank-sum test from sc.tl.rank_genes_groups function in Scanpy.

## Supporting information

Supplementary Section

Supplemental Datafile 1

Supplemental Datafile 2

## Data availability

Previously published ATAC-seq and cut&run-seq datasets are available from the GEO database under accession GSE266039. The merged scRNA-seq dataset, and newly generated cut&run-seq datasets will be uploaded to GEO. Proteomic data will be submitted to the PRIDE repository.

## Author contributions

Conceptualization: SM and PM. Investigation: all authors. Data curation and visualization: SM, AK, PM. Writing: original draft: SM and PM. Review and editing: All authors. Supervision and funding: PM.

## Acknowledgements

We thank Katia Georgopoulos and Toshimi Yoshida for sharing the *Chd4^Flox^* mice. We thank Zev Gartner, Chris McGinnis, David Cook, and Barbara Vanderhyden for generously providing Multi-seq protocols, reagents, and advice. We thank the Michael Dyer lab for providing datasets upon which this study relied. For scRNA-seq and cut&run-seq experiments, we also thank Katayoun Sheikheleslamy, Caroline Vergette, and Pearl Campbell from the *Stemcore Molecular Biology Core facility*, as well as Chris Porter from the *OHRI Bioinformatics Core Facility*. We thank Andrew Macklin from the *OHRI Proteomics Core Facility*. We thank Chloë van Oostende-Triplet and the *Cell Biology and Image Acquisition Core Facility*, and the staff of the uOttawa *Animal Care and Veterinary Service*. We thank members of the Mattar and Picketts labs for their ongoing support and input. This work was generously supported by the Canadian Institutes of Health Research (CIHR) Operating Grant (PJT-166032). Our Adnp research program is also generously supported by an Eagle’s Foundation for Autism Research grant: “Deciphering the role of chromatin remodelling complexes in ASD etiology”, awarded to P.M. and D.J.P. We additionally acknowledge the generous support of the New Frontiers in Research Fund (NFRFT-2022-00327), and from the Natural Sciences and Engineering Research Council of Canada (RGPIN-2026-05722). We also acknowledge the generous support of the Azrieli Research Program on Neurovascular Development (Azrieli Foundation). SS was generously supported by the David M. Shillito Scholarship in Ophthalmology Research. The project was also supported by an infrastructure grant from the Canada Foundation for Innovation (45420). PM gratefully holds the Gladys and Lorna J. Wood Chair for Research in Vision.

