## Supplementary Section for "Molecular dissection of the ChAHP complex subunits Adnp and Chd4 reveals common and divergent functions in retinal development"

Supplementary Figures

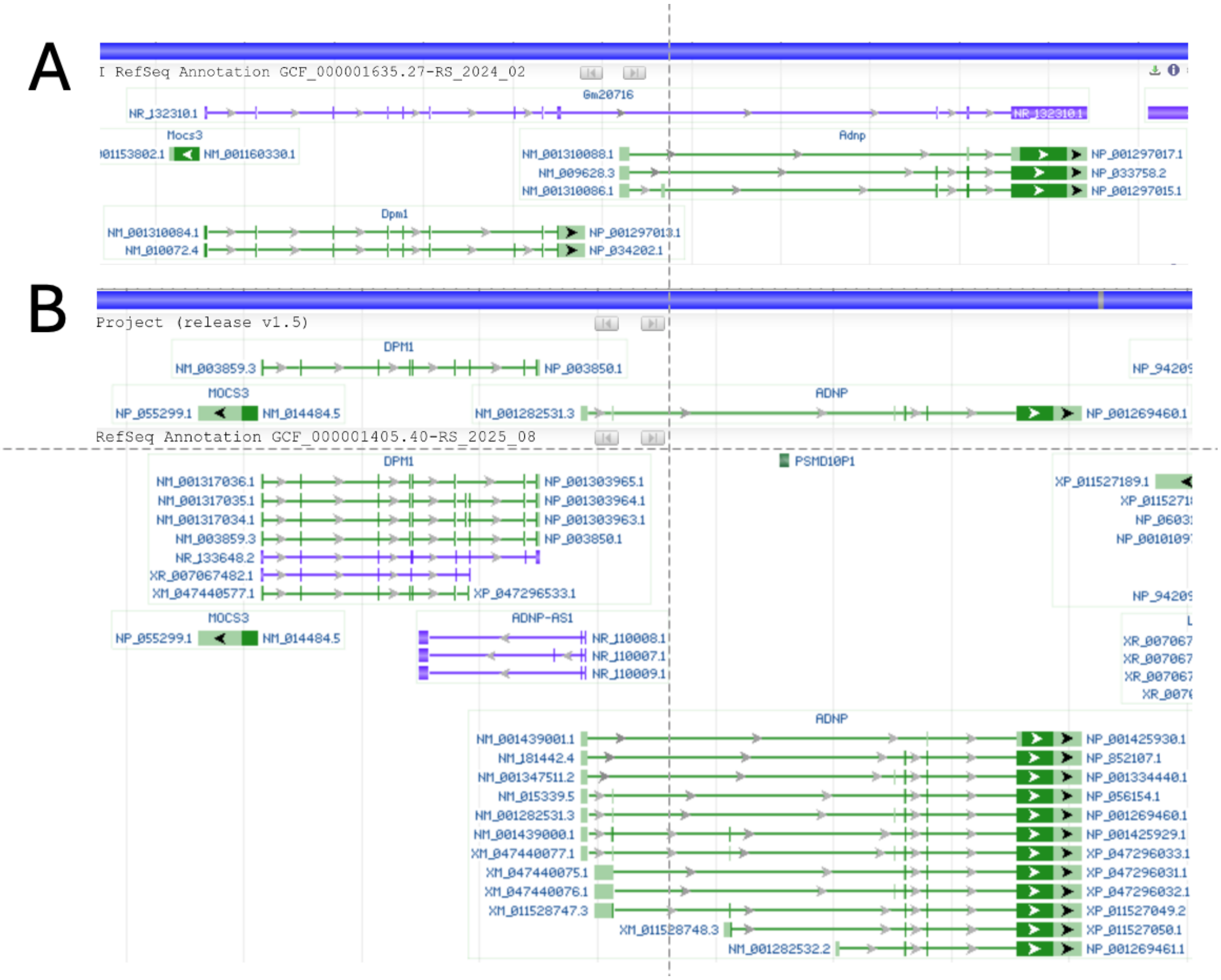


**Fig. S1. A *Dpm1-Adnp* readthrough transcript undermines murine RNA-seq profiling.** (A) Mouse mm11 genome. A readthrough transcript (NP_132310; purple) that splices *Adnp* exons to the upstream *Dpm1* gene double-annotates both sets of exons. (B) The readthrough transcript is absent from the human genome.

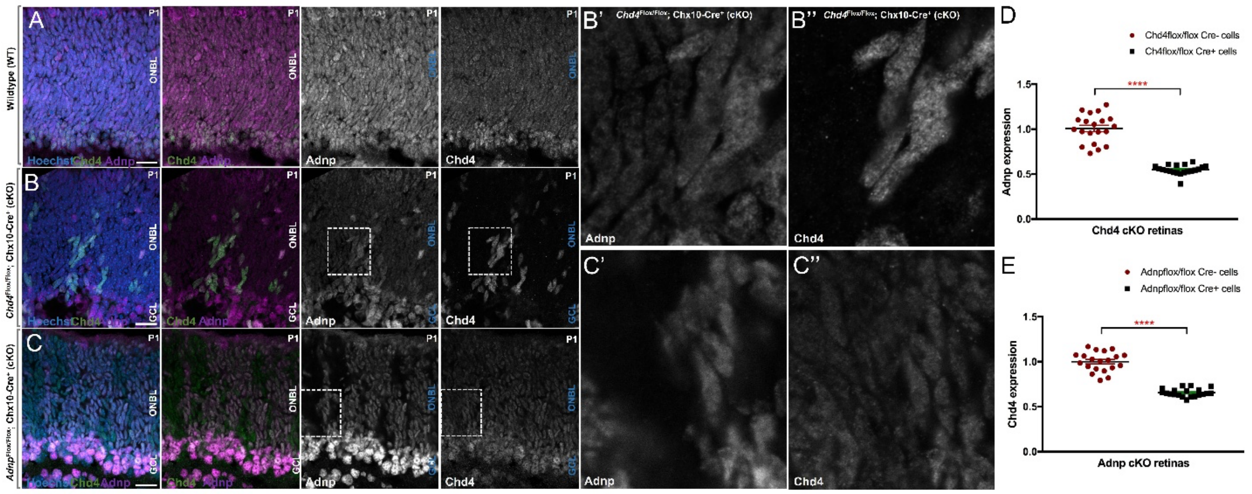


**Fig. S2 Genetic ablation reduces the levels of ChAHP partner proteins.** (A-C) Adnp and Chd4 co-staining performed on P1 retinal sections of wildtype (A), mosaic *Chd4* cKO (B) and mosaic *Adnp* cKO (C) retinas. Note that we rejected retinas with *Chx10-Cre* mosaicism in all other analyses. Adnp (magenta), Chd4 (green) and Hoechst (blue). Adnp and Chd4 alone are shown in grayscale. Dotted boxes indicate the magnified regions depicting Adnp (B') and Chd4 (B'') expression in *Chd4* cKO retina cells, or Adnp (C') and Chd4 (C'') expression in *Adnp* cKO retina cells. (D) Reduced Adnp expression in Cre positive (*Chd4* cKO) cells compared to Cre negative cells. (E) Reduced Chd4 expression in Cre positive (*Adnp* cKO) cells compared to Cre negative cells. Scale bar: 20 µm. ****$p < 0.0001$, via Student's t-test.

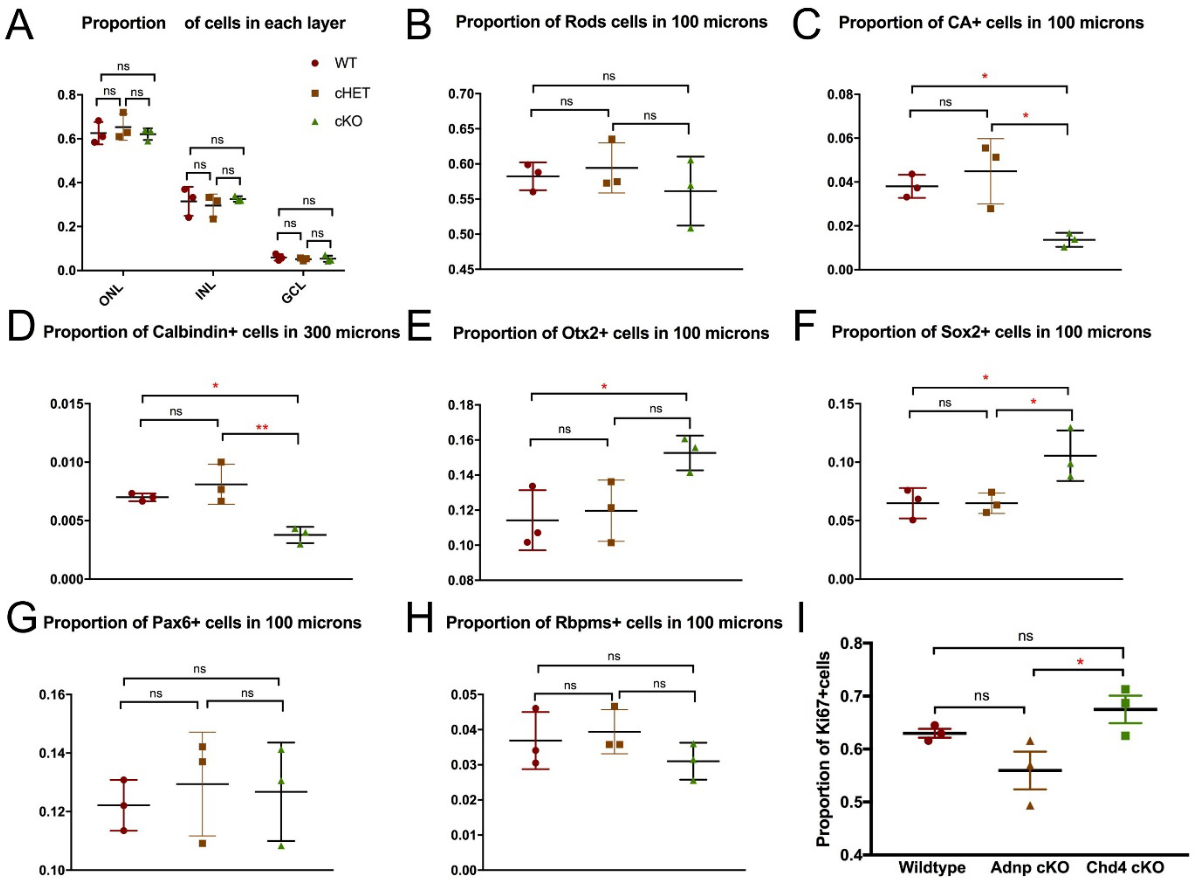


**Fig. S3. Proportional cell count data**. (A-H) P21 retinal counts. (A) Proportion of cells distributed among the 3 retinal layers. (B) Overall proportion of rods at P21. (C) Overall proportion of cone photoreceptors. (D) Overall proportion of horizontal cells. (E, F) Overall proportion of bipolar cells. (F) Overall proportion of Müller glia cells. (G) Overall proportion of amacrines. (H) Overall proportion of ganglion cells. (I) Overall proportion of Ki67 positive at P1. *$p<0.05$, **$p < 0.01$, via one-way ANOVA with the Tukey/Kramer post-hoc test.

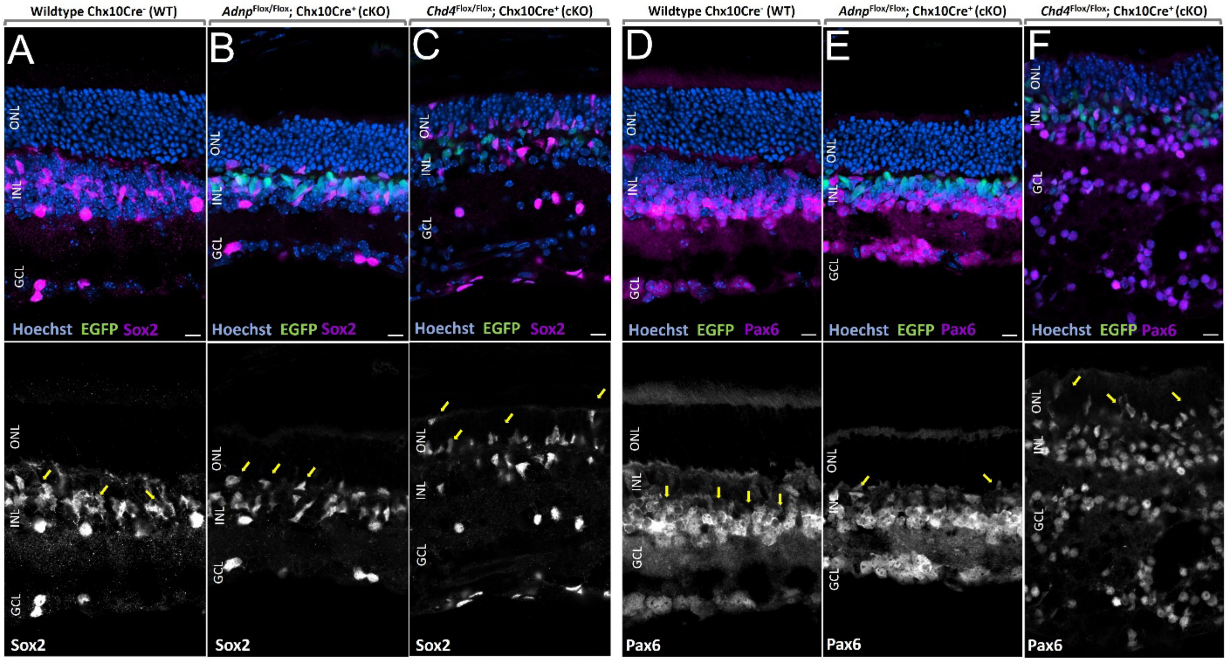


**Fig. S4. Ectopic cells in *Adnp* and *Chd4* cKOs.** (A-C) Sox2 staining performed on P21 retinal sections of wildtype, *Adnp* cKO and *Chd4* cKO. Sox2 (magenta) co-stained with Hoechst (blue) and Cre-EGFP (green). Sox2 staining alone is shown in grayscale. (D-F) Pax6 staining performed on P21 retinal sections of wildtype, *Adnp* cKO and *Chd4* cKO. Pax6 (magenta) co-stained with Hoechst (blue) and Cre-EGFP (green). Pax6 staining alone is shown in grayscale. Arrows indicate ectopic cells (likely Pax6+ Müller glia).

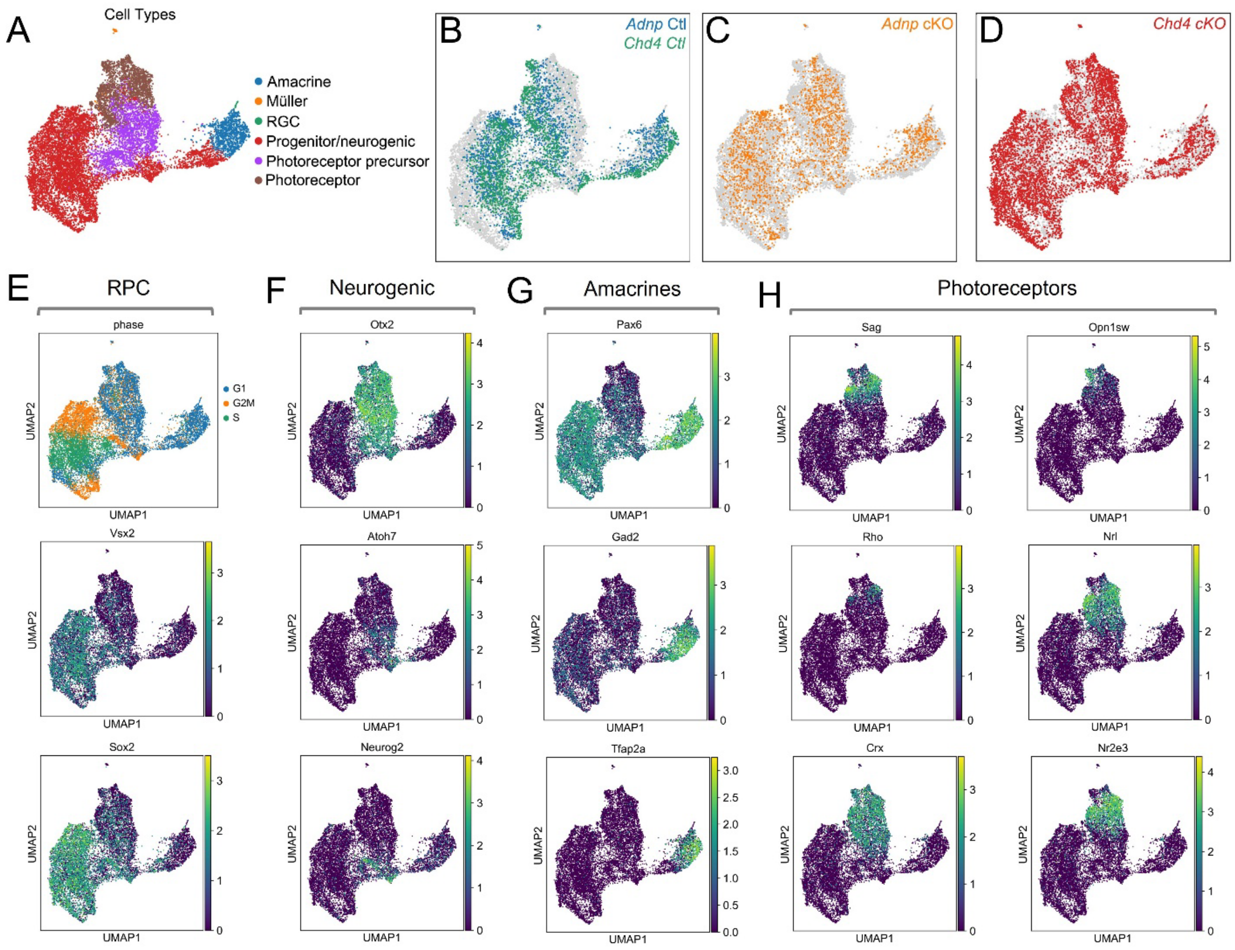


**Fig. S5. Cell type marker expression in the scRNA-seq dataset.** (A) UMAP clusters showing cell type annotation in the merged datasets. (B-D) UMAP projections of cells based on genotype. (E-I) UMAP projections of marker gene expression in the merged dataset. (E) RPCs identified based on cell-cycle phases and the expression of key progenitor-specific markers, such as *Vsx2* and *Sox2*. (F) Neurogenic cell markers such as *Otx2*, *Atoh7* and *Neurog2*. (G) Amacrine markers *Pax6*, *Gad2* and *Tfap2a*. (I) Photoreceptor markers *Sag*, *Rho*, *Crx*, *Opn1sw*, *Nrl*, and *Nr2e3*.

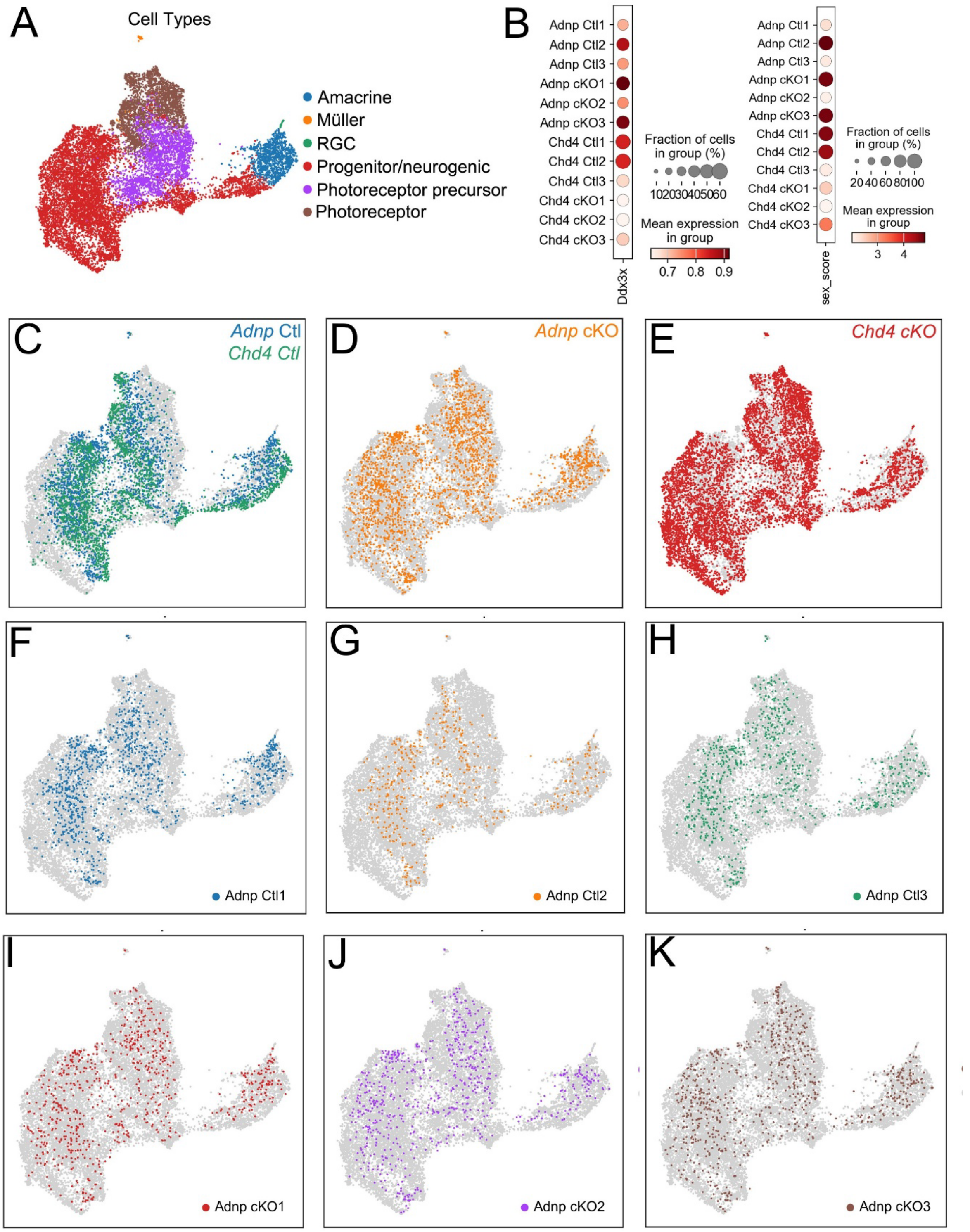


**Fig. S6. Additional analysis of the Adnp/Chd4 scRNA-seq dataset. (**A) Cell type annotation of UMAP clusters. (B) Inferred sex of the samples based on the ratio of reads mapped to sex chromosomes (chromosome X and Y genes). *Ddx3x* is only located on the X Chromosome. (C-

E) UMAP projection of cells separated based on genotype. (F-K) UMAP projection of cells separated by replicates of control (F-H) and *Adnp* cKO samples (I-K).

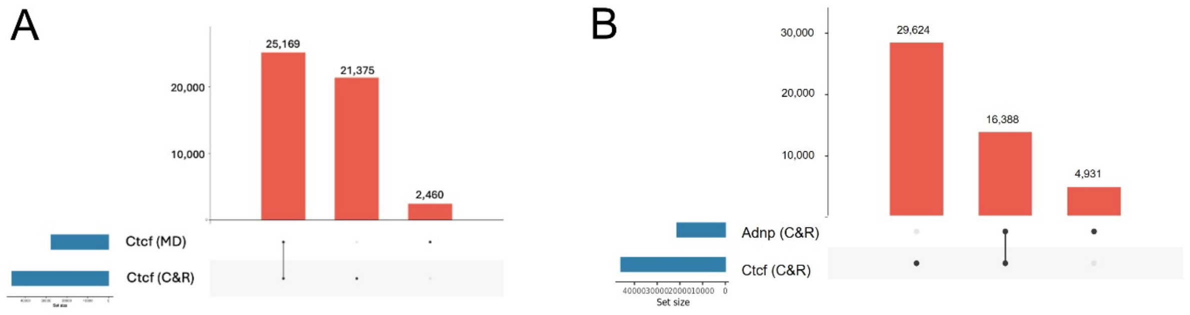


**Fig. S7.** (A) Benchmarking of Ctcf cut&run-seq data against published Ctcf ChIP-seq data from the Michael Dyer laboratory (MD) (Aldiri et al., 2017). (B) Comparison of Adnp and Ctcf peak numbers from retinal cut&run-seq.

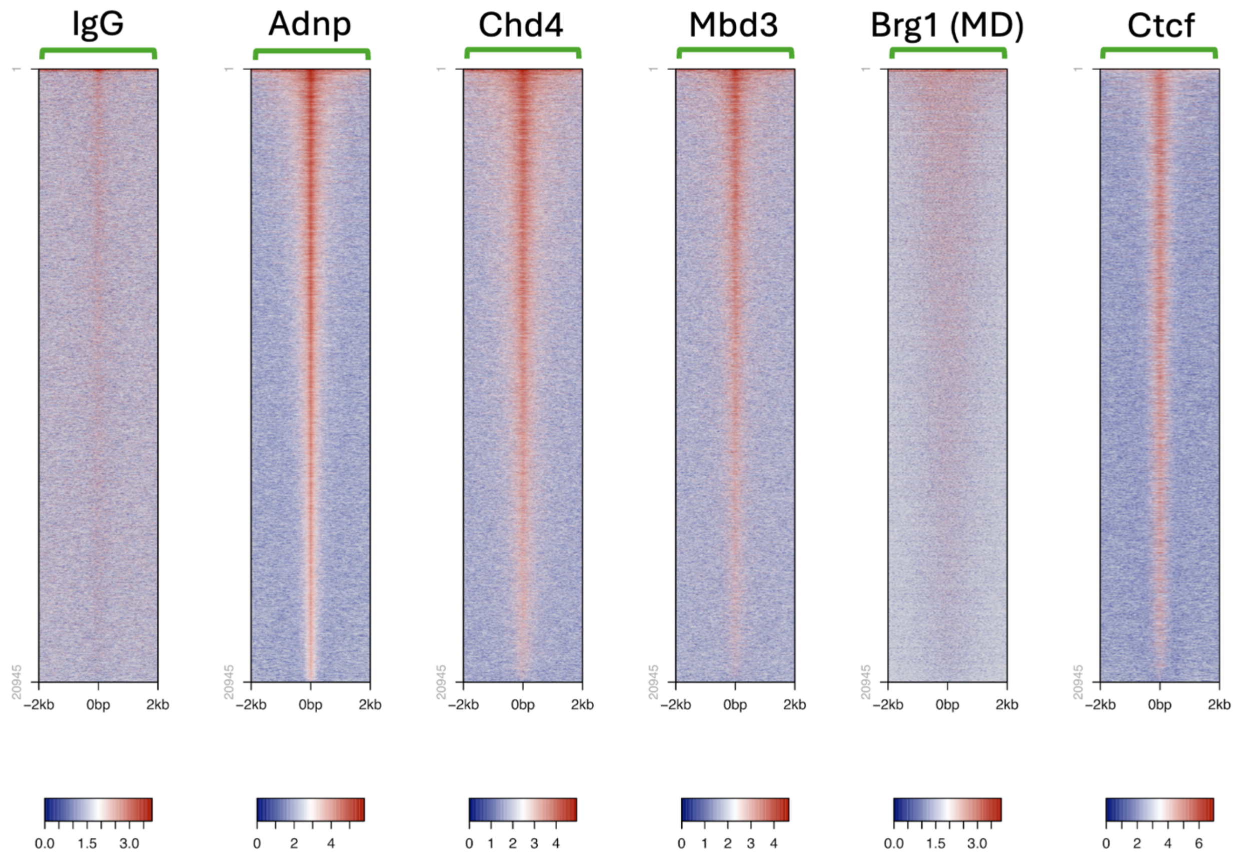


**Fig. S8. Absence of Brg1 at retinal Adnp peaks.** Cut&run-seq tracks compared against Brg1 ChIP-seq data generated by the Michael Dyer lab (MD) (Aldiri et al., 2015). Data are centered on called Adnp peaks.

**Table S1:** scRNA-seq cell type metrics

| Genotype | Cell Type | Cell number (post QC) |
|---|---|---|
| *Adnp* Ctl | (all) | 2331 |
| *Adnp* cKO | (all) | 2190 |
| *Chd4* Ctl | (all) | 2142 |
| *Chd4* cKO | (all) | 7438 |
| *Adnp* Ctl1 | (all) | 1094 |
| *Adnp* Ctl2 | (all) | 452 |
| *Adnp* Ctl3 | (all) | 785 |
| *Adnp* cKO1 | (all) | 730 |
| *Adnp* cKO2 | (all) | 687 |
| *Adnp* cKO3 | (all) | 773 |
| *Chd4* Ctl1 | (all) | 518 |
| *Chd4* Ctl2 | (all) | 457 |
| *Chd4* Ctl3 | (all) | 1167 |
| *Chd4* cKO1 | (all) | 2591 |
| *Chd4* cKO2 | (all) | 39 |
| *Chd4* cKO3 | (all) | 4808 |
| (all) | Progenitor/neurogenic | 7760 |
| (all) | Photoreceptor precursor | 2604 |
| (all) | Photoreceptor | 2524 |
| (all) | Amacrine cells | 1136 |
| (all) | Müller cells | 60 |
| (all) | Retinal ganglion cells | 17 |
| *Chd4* cKO | Progenitor/neurogenic | 4229 |
| *Chd4* cKO | Photoreceptor | 1434 |
| *Chd4* cKO | Photoreceptor precursor | 1339 |
| *Chd4* Ctl | Progenitor/neurogenic | 1230 |
| *Adnp* Ctl | Progenitor/neurogenic | 1189 |
| *Adnp* cKO | Progenitor/neurogenic | 1112 |
| *Adnp* Ctl | Photoreceptor precursor | 452 |
| *Adnp* cKO | Photoreceptor precursor | 449 |
| *Chd4* cKO | Amacrine cells | 401 |
| *Adnp* Ctl | Photoreceptor | 370 |
| *Chd4* Ctl | Photoreceptor precursor | 364 |
| *Adnp* cKO | Photoreceptor | 361 |
| *Chd4* Ctl | Photoreceptor | 359 |
| *Adnp* Ctl | Amacrine cells | 295 |
| *Adnp* cKO | Amacrine cells | 252 |
| *Chd4* Ctl | Amacrine cells | 188 |
| *Chd4* cKO | Müller cells | 26 |
| *Adnp* Ctl | Müller cells | 20 |
| *Adnp* cKO | Müller cells | 13 |
| *Chd4* cKO | Retinal ganglion cells | 9 |
| *Adnp* Ctl | Retinal ganglion cells | 5 |
| *Adnp* cKO | Retinal ganglion cells | 3 |
| *Chd4* Ctl | Müller cells | 1 |

**Table S2:** Oligonucleotide sequences. Multi-seq barcodes are highlighted in green.

| Name | Purpose | Sequence |
|---|---|---|
| Adnp-3'F | genotyping | 5′-TGGCGTAACAGTAAAGGAGAAGTAC-3′ |
| Adnp-3'R | genotyping | 5′-AGAGCTTCAGCATGACTTCCAGGTG-3′ |
| Mi-2β+F S | genotyping | 5′-CTCCAAGAAGAAGACGGCAGATCT-3′ |
| Mi-2 INR A | genotyping | 5′-GTCCTTCCAAGAAGAGCAAG-3′ |
| CRE-F | genotyping | 5′-AGGTGTAGAGAAGGCACTTAGC-3′ |
| CRE-R | genotyping | 5′-CTAATCGCCATCTTCCAGCAGG-3′ |
| Multiseq-1 | barcoding | 5′-CCTTGGCACCCGAGAATTCCAGGAGAAGAAAAAAAAAAAAAAAAAAAAAAAAAAAAAAA-3′ |
| Multiseq-2 | barcoding | 5′-CCTTGGCACCCGAGAATTCCACCACAATGAAAAAAAAAAAAAAAAAAAAAAAAAAAAAA-3′ |
| Multiseq-3 | barcoding | 5′-CCTTGGCACCCGAGAATTCCATGAGACCTAAAAAAAAAAAAAAAAAAAAAAAAAAAAAA-3′ |
| Multiseq-4 | barcoding | 5′-CCTTGGCACCCGAGAATTCCAGCACACGCAAAAAAAAAAAAAAAAAAAAAAAAAAAAAA-3′ |
| Multiseq-5 | barcoding | 5′-CCTTGGCACCCGAGAATTCCAAGAGAGAGAAAAAAAAAAAAAAAAAAAAAAAAAAAAAA-3′ |
| Multiseq-6 | barcoding | 5′-CCTTGGCACCCGAGAATTCCATCACAGCAAAAAAAAAAAAAAAAAAAAAAAAAAAAAAA-3′ |
| Multiseq-7 | barcoding | 5′-CCTTGGCACCCGAGAATTCCAGAAAAGGGAAAAAAAAAAAAAAAAAAAAAAAAAAAAAA-3′ |
| Multiseq-8 | barcoding | 5′-CCTTGGCACCCGAGAATTCCACTATAGTAAAAAAAAAAAAAAAAAAAAAAAAAAAAAAA-3′ |
| Multiseq-9 | barcoding | 5′-CCTTGGCACCCGAGAATTCCATAAAATCCAAAAAAAAAAAAAAAAAAAAAAAAAAAAAA-3′ |
| Multiseq-10 | barcoding | 5′-CCTTGGCACCCGAGAATTCCAGTATATGTAAAAAAAAAAAAAAAAAAAAAAAAAAAAAA-3′ |
| Multiseq-11 | barcoding | 5′-CCTTGGCACCCGAGAATTCCATAATCAACAAAAAAAAAAAAAAAAAAAAAAAAAAAAAA-3′ |
| Multiseq-12 | barcoding | 5′-CCTTGGCACCCGAGAATTCCAGTAGCACTAAAAAAAAAAAAAAAAAAAAAAAAAAAAAA-3′ |

**Table S3:** Antibodies and dilutions.

| Antigen | Species | Dilution | Supplier | Catalog | RRID | Application |
|---|---|---|---|---|---|---|
| Adnp | Goat | 1:200<br>1:1000<br>1:100 | R&D Systems | AF5919 | AB_1964493<br>(discontinued) | IHC,<br>IP/Western<br>cut&run |
| Chd4 | Rat | 1:200 | Biolegend | 942302 | AB_2888898 | IHC |
| Chd4 | Rabbit | 1:1000<br>1:100 | Abcam | ab72418 | AB_1268107 | Western,<br>cut&run |
| Cleaved Caspase-3 (Asp175) | Rabbit | 1:500 | Cell Signaling | 9579 | AB_10897512 | IHC |
| Cone arrestin | Rabbit | 1:200 | Millipore Sigma | AB15282 | AB_1163387 | IHC |
| Ctcf | Rabbit | 1:100 | Cell Signaling | 3418 | AB_2086791 | cut&run |
| Ki67 | Rabbit | 1:100 | Millipore Sigma | SAB5500134 | AB_2892217 | IHC |
| Mbd3 | Rabbit | 1:100 | Abcam | EPR18258 | Not found | cut&run |
| Otx2 | Goat | 1:500 | R&D Systems | AF1979 | AB_2157172 | IHC |
| Pax6 | Rabbit | 1:500 | Novus | NBP2-19711 | AB_3264565 | IHC |
| Pax6 | Rabbit | 1:500 | Proteintech | 12323-1-AP | AB_2159695 | IHC |
| Rbpms | Guinea pig | 1:200 | Millipore Sigma | ABN1376 | AB_2687403 | IHC |
| Sox2 | Goat | 1:500 | R&D Systems | AF2018-SP | AB_355110 | IHC |
| anti-guinea pig Alexa Fluor™ 555 | Goat | 1:1000 | Rockland | 606-142-129 | AB_1961616 | 2º (IHC) |
| anti-goat Alexa Fluor™ 488 | Donkey | 1:1000 | Jackson | 705-547-003 | AB_2340431 | 2º (IHC) |
| anti-goat DyLight™ 650 | Donkey | 1:1000 | Novus | NBP1-75604 | AB_11018252 | 2º (IHC) |
| anti-mouse Alexa Fluor™ 555 | Donkey | 1:1000 | Invitrogen | A-31570 | AB_2536180 | 2º (IHC) |
| anti-mouse Alexa Fluor™ 647 | Donkey | 1:1000 | Invitrogen | A-31571 | AB_162542 | 2º (IHC) |
| anti-rabbit Alexa Fluor™ 488 | Donkey | 1:1000 | Jackson | 111-545-003 | AB_2338046 | 2º (IHC) |
| anti-rabbit Alexa Fluor™ 647 | Donkey | 1:1000 | Jackson | 711-607-003 | AB_2340626 | 2º (IHC) |
| anti-rat DyLight™ 550 | Donkey | 1:1000 | Invitrogen | SA510027 | AB_2556607 | 2º (IHC) |
| anti-Rabbit HRP | Donkey | 1:10 000 | GE Healthcare | NA934 | AB_772206 | 2º (Western) |
